# Therapeutic Monoclonal Antibodies Repurposing in Oncology via IMGT/mAb-KG Embeddings

**DOI:** 10.1101/2025.09.05.674444

**Authors:** Gaoussou Sanou, Taciana Manso, Konstantin Todorov, Véronique Giudicelli, Patrice Duroux, Sofia Kossida

**Affiliations:** IMGT, The International ImMunoGeneTics Information System, Institute of Human Genetics, National Center for Scientific Research/University of Montpellier, 141 Rue de la Cardonille, Montpellier, 34090, France; Montpellier Laboratory for Computer Science, Robotics and Microelectronics, National Center for Scientific Research/University of Montpellier, 161 Rue Ada, Montpellier, 34095, France; University Institute of France, 1 rue Descartes, Paris, 75231, France

**Keywords:** Immunogenetics, Oncology, Monoclonal antibody, Knowledge Graph, Knowledge Graph embeddings, Drug repurposing

## Abstract

**Background:** Cancer remains one of the leading causes of mortality world-wide, accounting for approximately 9.7 million deaths in 2022. Faced with this significant public health challenge, therapeutic monoclonal antibodies (mAbs) have emerged as promising alternatives that may minimize the side effects associated with conventional treatments such as radiotherapy and chemotherapy. To support mAb research and development, IMGT®, the international ImMunoGeneTics information system, has established two standardized data sources namely IMGT/mAb-DB, a comprehensive database for mAbs, and, more recently, IMGT/mAb-KG, a dedicated knowledge graph for mAbs. Despite these advances, the development of therapeutic mAbs remains both time-consuming and financially burdensome—costs can reach up to $2.8 billion. To address this challenge and accelerate cancer treatment, mAb repurposing represents a promising alternative.

**Results:** In this study, we leveraged a subset of IMGT/mAb-KG, dedicated to the oncology domain, to develop a scientific hypothesis generation application for mAb repurposing. This application, based on knowledge graph embedding techniques, is designed to suggest potential mAb candidates for novel oncology applications. A user-friendly web interface provides access to the tool, incorporating visual support to facilitate the interpretation of generated hypotheses. This application is a decision support tool aiming to accelerate the discovery of new therapeutic applications for existing mAbs.

**Conclusion:** Our application demonstrates the potential of knowledge graph embedding techniques in the oncology domain by enabling the repurposing of existing mAbs for new therapeutic uses. Using this tool, we have identified two novel mAbs, loncastuximab tesirine and glofitamab, both currently undergoing clinical trials for the treatment of chronic lymphocytic leukemia. This decision-support tool thus facilitates the discovery of new therapeutic opportunities by effectively repositioning existing mAbs for oncological indications, potentially accelerating the development of cancer therapies and addressing critical public health needs.

## 1 Introduction

Cancers are one of the leading causes of mortality worldwide. In fact, in 2022, 20 million new cases of cancer were detected with more than 9.7 million deaths from cancer in the world [1]. According to the World Health Organization (WHO), cancer is the second leading cause of death before the age of 70 years [2]. Our immune system plays an intrinsic role in the physiological fight against cancers, including their detection and elimination. Consequently, understanding the basic principles of cancer-immune system interactions allows us to develop therapeutic strategies to activate and strengthen the immune system for cancer treatment. Hence, therapeutic monoclonal antibodies (mAbs) have been developed.

mAbs are laboratory-engineered molecules designed to function as surrogate antibodies, capable of restoring, enhancing, modifying, or mimicking the immune system’s targeting of undesired cells, such as cancerous cells. In cancer’s therapy, mAbs have the potential to specifically target, recognize and trigger the destruction of cancer cells, avoiding side effects stemming from chemotherapy or radiotherapy [3, 4]. In addition to the cancer therapy, mAbs are used to treat many other diseases such as infectious diseases, cardiovascular diseases, hematological diseases, autoimmune diseases, etc.

Nowadays, the mAbs represent one of the principal biological drugs in development and clinical trials in the pharmaceutical industries [5]. Consequently, the mAbs market has exploded these last years. In fact, the mAbs market was $75 billions in 2013 [6], $150 billions in 2019 and is expected to reach $300 billions by 2025 [7]. Given the interest in mAbs and their vast medical capabilities, several standardized data sources, such as the [National Cancer Institute Thesaurus (NCIt)](https://ncithesaurus.nci.nih.gov/)[8], [Thera-SAbDab](https://opig.stats.ox.ac.uk/webapps/sabdab-sabpred/sabdab) [9] or PladDab [10] have emerged. The goal of these data sources is to provide standardized information on mAbs to facilitate their comprehension and accelerate their development process. To support these efforts, we developed IMGT/mAb-DB a database dedicated to mAbs and therapeutic proteins [11] and lately IMGT/mAb-KG, a knowledge graph (a data representation framework that describes real-world entities and their interrelations in a graph.) integrating data from IMGT/mAb-DB [12]. To date, IMGT/mAb-KG provides access to 1,586 mAbs targeting 500 targets in over 500 clinical indications and offers detailed insights into their mechanisms of action, design, and associated studies [12]. In addition to the use of standardized and integrated data sources such as IMGT/mAb-KG, a crucial factor in accelerating the development process of mAbs is the ability to discover and validate novel therapeutic molecules or targets for both existing and new mAbs [4].

Recently, the discovery of new targets has significantly slowed [13]. The introduction of new drugs including mAbs to the market is becoming increasingly challenging, largely due to difficulties in identifying novel active compounds and the increasing complexity of their mechanisms of action [13, 14]. Furthermore, the drug development process, from initial conception to market approval, typically spans 13–15 years and has a success rate of less than 10% [13–16]. This process is not only time-consuming and labor-intensive, but also financially burdensome, with the average cost per drug estimated at $2.8 billion for pharmaceutical companies [14, 17].

Given these increasing challenges, alternative strategies, particularly drug repurposing, which involves utilizing already-approved drugs for new therapies, have gained increasing attention. By leveraging existing knowledge of these approved compounds, drug repurposing aims to discover new therapeutic applications, often taking advantage of the “off-target” phenomenon (a drug initially designed for a specific target that interacts with additional unintended targets) or the pathway overlap and the mechanism of action (the mAb’s target can be involved in different diseases)[13, 15]. During the 2019 global health crisis, drug repurposing was widely employed to identify existing treatments for COVID-19 [18]. For example, antiviral drugs such as baloxavir, azvudine, and darunavir have been to combat COVID-19 [18], whereas baricitinib, which was originally used for rheumatoid arthritis, has been utilized to treat COVID-19-induced bilateral pneumonia [15]. In oncology, denosumab (PROLIA®), an anti-TNFSF11 antibody, was initially investigated for osteoporosis [19] and received FDA approval in 2010. It is now being repurposed for breast cancer prevention for women, with BRCA1 mutations, because of its ability to inhibit TNFSF11 involved in breast cancer initiation and progression [20].

There are two main approaches to drug repurposing: the first is based on biological experimentation, which is both costly and time-consuming, while the second leverages computational methods to predict potential new active molecules, which are then validated through biological assays. Among these computational approaches, approaches based on virtual screening and knowledge graphs (KG) have garnered attention in recent years. The virtual screening approach consists of using computational tools to predict potentially bioactive compounds from multiple libraries of small molecules [21]. The idea of virtual screening is to predict how well the structure of a drug molecule fits or interacts with the structure of a target protein. However, this approach is subject to a high rate of false positives. In fact, highlighting, many of the drug-target protein interactions predicted *in vitro* or *in vivo* experiments is difficult [22]. The second approach, which is based on the KG, considers the complex relationships among diverse biomedical data related to a drug and its targets beyond the structural aspect of the drug-target protein interaction. Indeed, several studies have utilized KG to facilitate drug repurposing, with promising results [18, 23, 24].

The approaches based on the KG for drug repurposing are formulated as a prediction task where, for a given drug, we predict the potential clinical indications or for a given clinical indication, we predict the potential mAb candidates that can be used. In practice, we apply representation learning techniques on KG to embed them into numerical vectors. This approach ensures that the embeddings of similar entities within the KG are clustered together in the resulting vector space [25]. These embeddings are then processed by machine learning tools to predict drug candidates for clinical indications.

In this work, we took advantage of the KG and KG embedding (KGE) techniques to build an intelligent application, generating new scientific hypotheses regarding the potential mAbs candidates for a given clinical indication in the oncology domain. For that, we extracted a subset of IMGT/mAb-KG, named IMGT/mAbOnco-KG, dedicated to the oncology domain, then applied representation learning techniques to generate embedding vectors of IMGT/mAbOnco-KG. We evaluated 28 KG embedding models, and then selected the top ten best models for a second test to optimize their performances. Once optimized, we evaluated the models on the mAbs and clinical indications relations. To provide access to this application, we extended IMGT/mAb-KG interface by adding a new section.

The remainder of the article is organized as follows. In section 2, we provide a background of this work including the definition of KG, their embeddings, and applications, and finish this section, by presenting IMGT/mAb-KG. Section 3 presents the dataset and the setup of the different experiments carried out in this work, including the link prediction task formulation, the evaluation, and the experiment settings. Section 4 presents the different results of the subsequent experiments and the scientific hypothesis generation application built for mAb candidates generation. Section 5, provides use cases of the hypothesis generation application in ovarian cancers and chronic lymphocytic leukemia. Finally, we conclude and outline directions for future work.

## 2 Background

### 2.1 Knowledge Graphs

A KG is a data representation framework that describes real-world entities and their interrelations in a graph. The entities are the nodes of the graph and the relations between entities are the edges. In a KG, 𝒢, a fact ℱ is described with a triplet (*h, r, t*) where *h* is the head entity and *t* is the tail entity linked to the head with the relation *r*. Consequently, a KG can be described by 𝒢 = {ℰ, ℛ, ℱ} with *h, t* ∈ ℰ, *r* ∈ ℛ and (*h, r, t*) ∈ ℱ.

KG facilitated the integration and federation of diverse data sources by unifying their content. These last years, we assisted in the emergence of KG in the biomedical domain integrating data from diverse sources including genomic, transcriptomic, proteomic, clinical trial or pharmacogenomic data [26]. In fact, different KG have emerged for different purposes in the biomedical domain, for instance the Clinical Trials Knowledge Graph ([CTKG](https://github.com/ninglab/CTKG)) is dedicated to clinical trials in drug studies [27]. In the context of the world pandemic, COVID-19, KG-COVID-19 was created to accelerate research on coronavirus [28]. For clinical purposes, a personalised medicine KG named Clinical Knowledge Graph ([CKG](https://ckg.readthedocs.io/en/latest/INTRO.html)) was developed. Aiming to facilitate access to immunogenetics information inside IMGT, the IMGT-KG, the first knowledge graph in immunogenetics has seen the light of day [29] and lately, IMGT/mAb-KG dedicated to mAbs and other proteins for therapeutic application was created [12].

### 2.2 Knowledge Graph Embeddings and Applications

The integration of biomedical data in a KG opens up new perspectives for data exploration as well as for the development of intelligent applications and the generation of new hypotheses of links between biomedical entities via machine learning [15, 30]. These applications allow us to understand the complexity of biological relations and hence discover new interactions. For example, in cancer therapies, we can leverage these applications to find new targets for existing drugs, leading to the repurposing of existing drugs for new clinical indications [31].

To implement these applications, one technique consists of representing the KG in a vector space of dimension *d*, such that the entities that are semantically and structurally close in the KG are grouped together in the vector space: this is representation learning or KG embedding in the case of KG [25]. The vector representations or embeddings capture the semantic relations between different entities in the KG. For instance, in the vector space, mAbs will be clustered and distinguished from targets, which will also be clustered. In general, KGE are generated under a KG completion or link prediction goal, this task involves predicting the missing entity or relation in a triple: (?, *r, t*), (*h*, ?, *t*), or (*h, r*, ?). In fact, the idea is to predict whether a new fact ℱ (*h, r, t*) ∉ 𝒢 in the KG is likely to be true, so it can be added to the KG as a new fact [32]. For that, KGE models define a score function *f* (*h, r, t*) that measures the plausibility of each fact (*h, r, t*) in 𝒢. The objective of the models is to choose f such that the score *f* (*h, r, t*) of a correct triple (*h, r, t*) is higher than the score *f* (*h*^*′*^, *r*^*′*^, *t*^*′*^) of an incorrect triple (*h*^*′*^, *r*^*′*^, *t*^*′*^) [32].

Figure 1, provides a summary of the KGE training process:

**Fig. 1.**
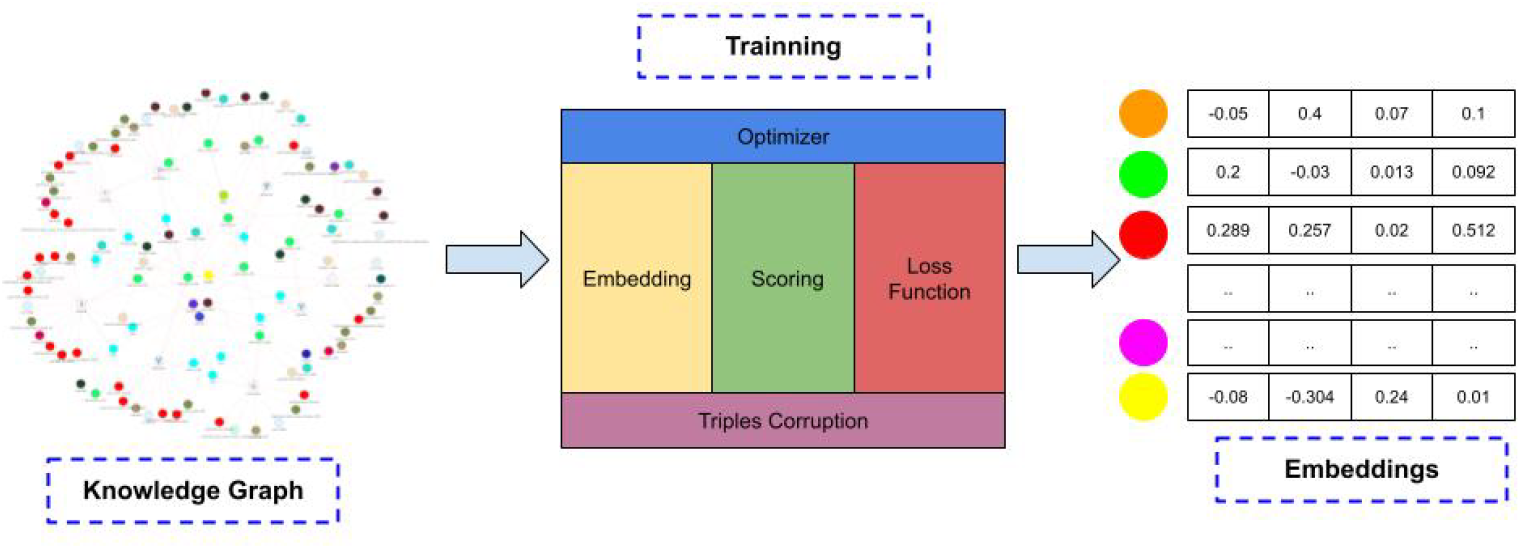
From KG to KG embeddings: KGE model training

- First, we randomly initialize the vectors corresponding to the representations of entities and relations, via either a uniform distribution or a Gaussian distribution [33].
- Then, we generate negative examples for each triplet existing in the KG. Thus, for a triplet *x*, (*h, r, t*) of the KG, we will corrupt it, by replacing either *h, r* or *t* such that we obtain a triplet *x*^*′*^ that does not exist in the KG [34]. Consequently, we will have a graph representing the corrupted triplets 𝒢*′* = {ℰ, ℛ, ℱ*′*} defined by:

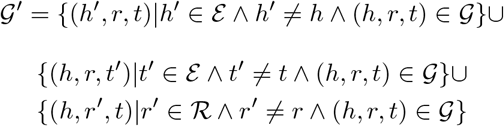
- Once the corrupted triplets are generated, a score function is used to assign a score to the positive and negative triplets *f* (*x*) and *f* (*x*^*′*^) [33, 34]. This score function is the key component in the KGE models, and the assigned score will be maximized for the observed triplets (positive: *f* (*x*)) in the KG and minimized for the unobserved triplets (negative: *f* (*x*^*′*^)).
- To maximize the score of the positive triplets, a loss function is used to compare the two scores *f* (*x*) and *f* (*x*^*′*^). Thus, during the optimization step, the model learns the representations of the entities and relations by adjusting the values of their vectors in such a way as to reduce the loss, which implies maximizing the score of the positive triplets.
- Once the loss is sufficiently reduced, we obtain representations of the entities and relations, thus capturing the semantics of the KG.

Based on the definition of the score function, different KGE models have been proposed, including translational models (TransE, TransR, TransH, TransD, PairE, etc), tensor-based models (RESCAL, DistMult, HolE, etc), complex vector-based models (ComplEx, QuatE, RotatE, etc) or graph neural network-based models (R-GCN, CompGNN, etc).

Once we apply KGE models to learn the KG representation, we obtain rich numerical vectors that capture the semantic and the structure of the KG. These vectors can be used in downstream tasks (Figure 2), including classification (for a given entity, predict it belonging class), link prediction (predict new links between existing entities) or clustering and visualization (group similar entities and visualize them).

**Fig. 2.**
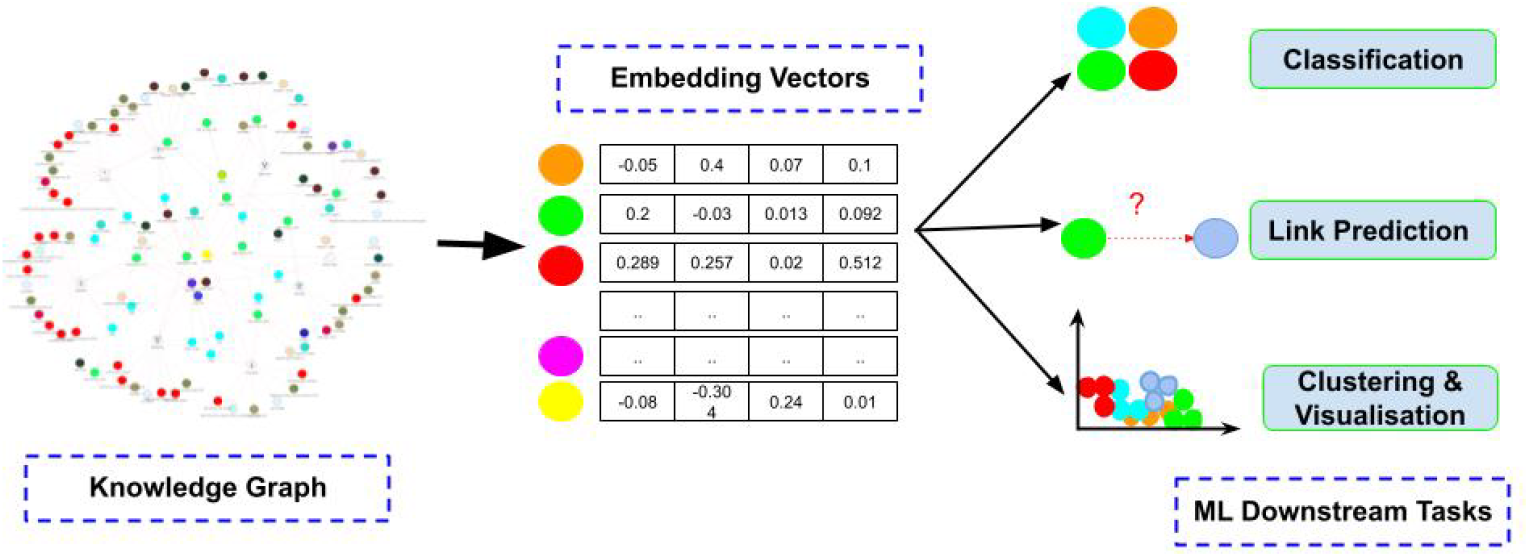
Downstream tasks using KGE vectors

### 2.3 IMGT/mAb-KG: the Knowledge Graph for Therapeutic Monoclonal Antibodies

Recently, in [12], we introduced IMGT/mAb-KG, the IMGT-KG for therapeutic monoclonal antibodies. IMGT/mAb-KG integrates data from the IMGT/mAb-DB, a unique expertized resource on mAbs [35], with related data in IMGT-KG including genomics and proteomics information. To date, IMGT/mAb-KG provides access to 149,354 triplets with 115 relations linking 22,620 entities. In addition, IMGT/mAb-KG is linked to other resources such as [Thera-SAbDab](https://opig.stats.ox.ac.uk/webapps/sabdab-sabpred/therasabdab/search/), [PharmGKB](https://www.pharmgkb.org/), [PubMed](https://pubmed.ncbi.nlm.nih.gov/) or [HGNC](https://www.genenames.org/), making it an indispensable resource in the research and development of mAbs.

Figure 3 shows the data model of IMGT/mAb-KG, an exhaustive description can be found in [12]. In IMGT/mAb-KG, a mAb is identified as a pharmacological substance with (`sio:SIO_0000291`) at least one target in a given taxon. It can have an origin cellular clone and can be associated with biosimilar products. Furthermore, a mAb can be linked to an access number inside IMGT-KG, thus bridging the two KG.

**Fig. 3.**
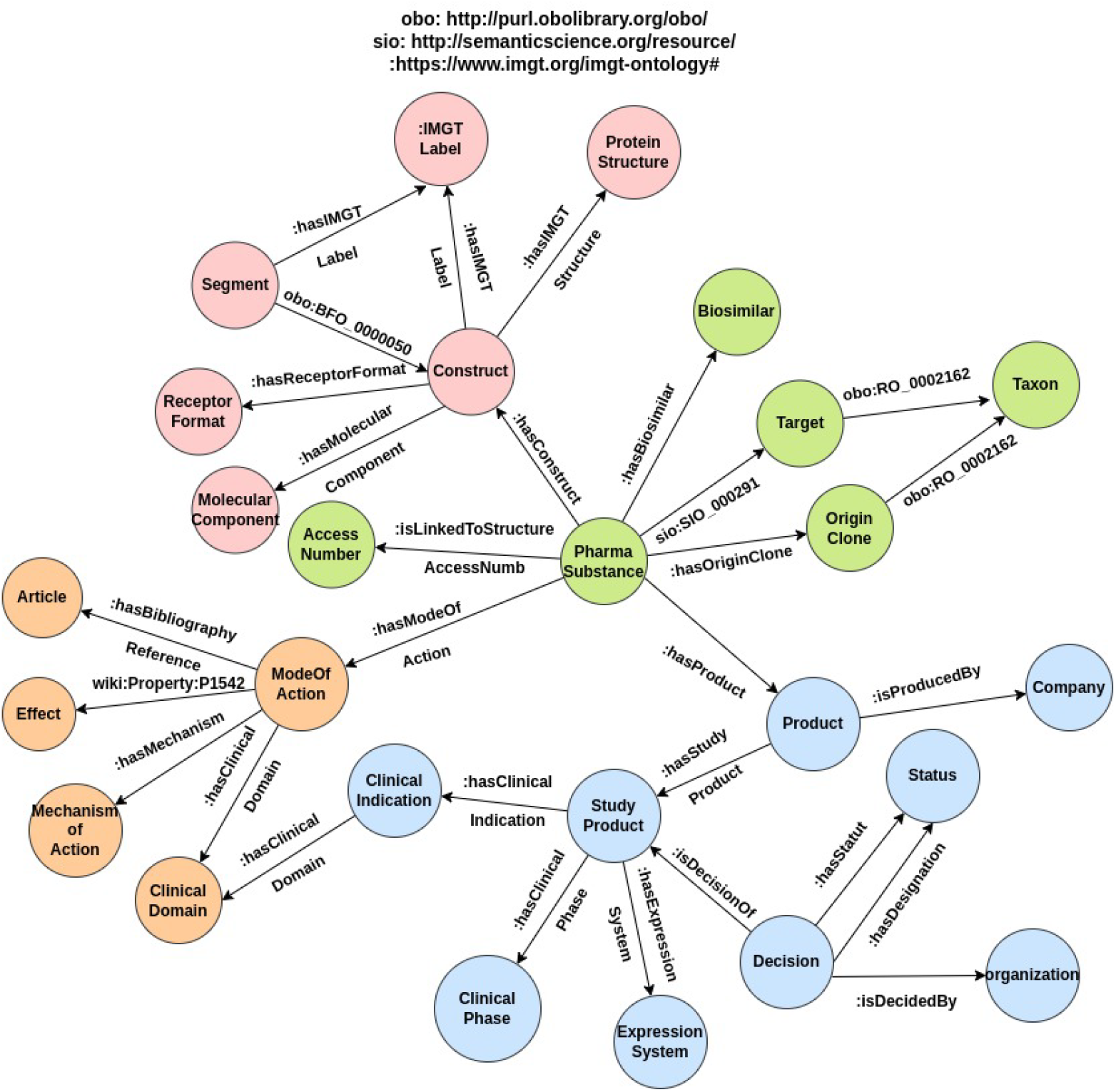
IMGT/mAb-KG data model. The green color represents the mAb level, the pink color represents the construct level, the blue color describes the product level and the orange color represents the mode of action level. For readable purposes, we replace the entity identifiers with their label (human-readable text) and we keep relation identifiers with the identifier. More details can be found in [12] To ensure clarity, we indicate the source name (namespace) before the relation when it is imported from an external source, for instance obo: http://purl.obolibrary.org/obo/ and sio: http://semanticscience.org/resource/.

A mAb is made with a construct describing its constitution with a receptor format and a molecular component. The construct is labelled with a unique IMGT label and consists of at least one segment with also an IMGT label. In reverse, the segment is a part of (`obo:BFO_0000050`) a construct. Moreover, a construct might be associated with a protein structure inside IMGT-KG.

A mAb is associated with at least one product, describing the clinical studies related to the mAb. A product is the result of a company’s production and invariably undergoes at least one evaluation or study. Each study focuses on a distinct clinical indication, primarily diseases, with a clinical phase and can be expressed within an expression system. The study product caan be subject to a decision taken by an organization.

Finally, the mAb can have a mechanism of action (MOA) describing how the mAb acts in the clinical indication. The MOA defines the mechanism of the mAb, detailing its effects by the has effect (`wiki:Property:P1542`) relation, and is linked to the bibliography information and a clinical domain.

## 3 Experiment Setups

### 3.1 Dataset

In our study, we consider a subset of IMGT/mAb-KG dedicated only to mAbs with applications in oncology. In fact, we created a sub-graph, named IMGT/mAbOnco-KG, that describes mAbs and all the related information but restricted it to the oncology domain. In this new dataset, we count 684 mAbs resulting from 152 different receptor formats, which were tested in 1812 studies on 228 targets in 247 oncology indications (Figure 4).

**Fig. 4.**
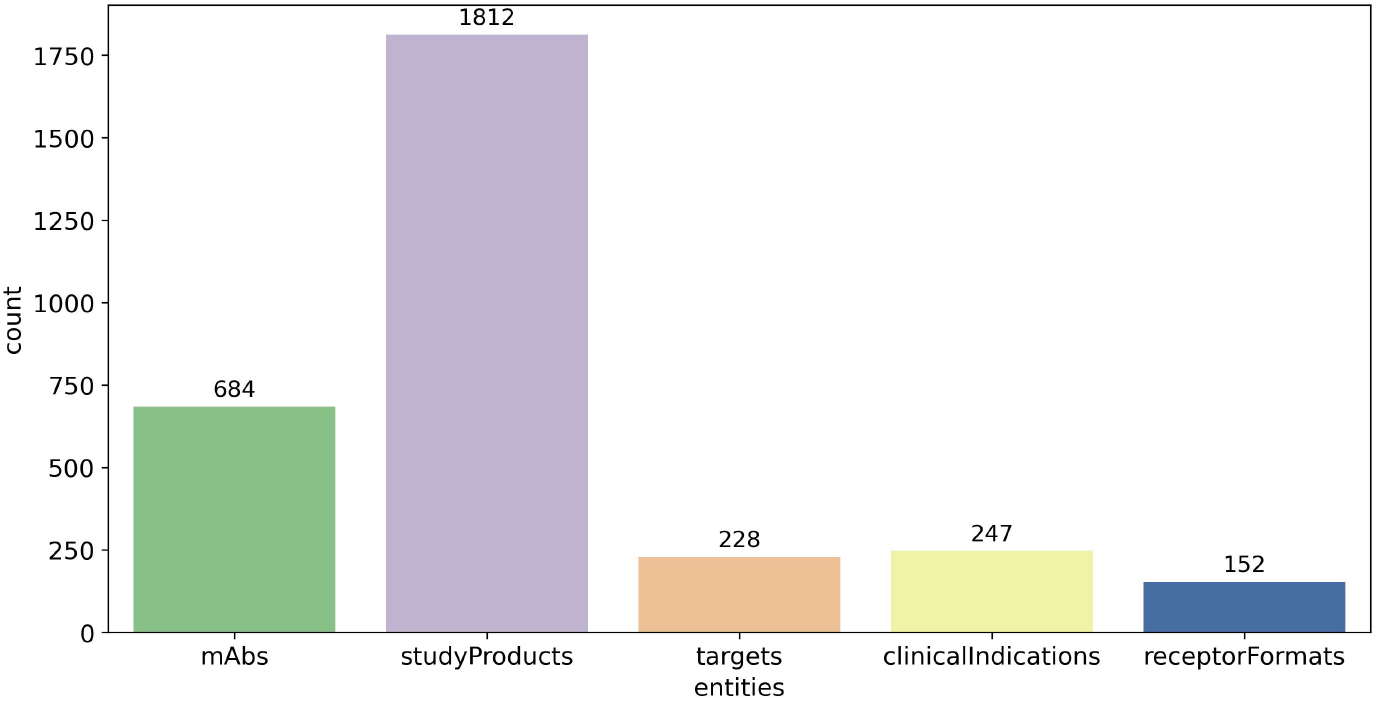
Some entities distributions in the dataset: IMGT/mAbOnco-KG

In the dataset, many studies have been conducted on solid tumour (264) and multiple myeloma (216). In the case of solid tumour, the protein that is most targeted is the PDCD1 and the protein TNFRSF17 in the case of multiple myeloma (Figure 5). Figure 5 shows the top ten clinical indications studied in the dataset and the related targets involved in these indications.

**Fig. 5.**
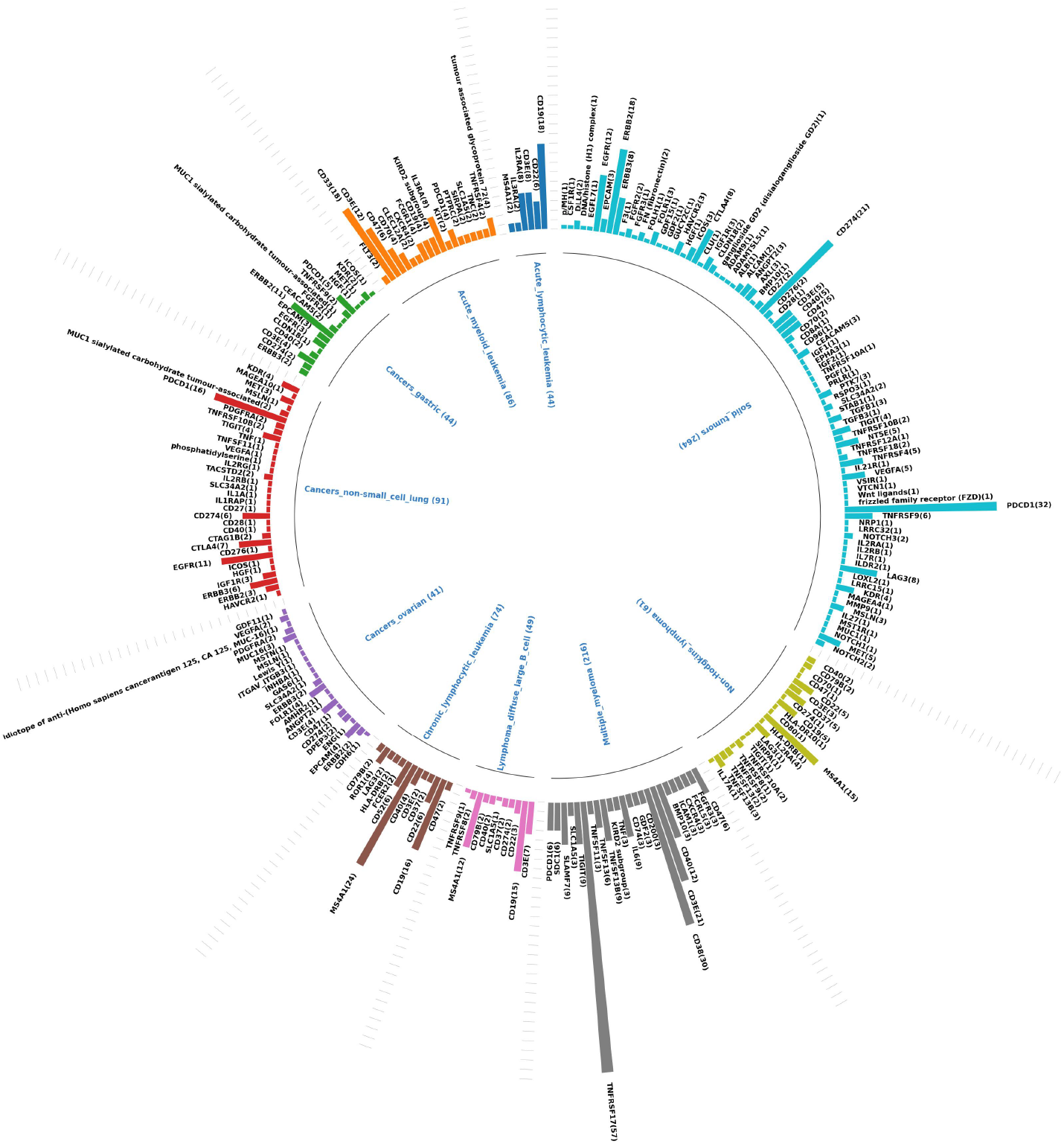
Chord diagram of the top ten clinical indications studied in the dataset and their target distributions

To generate embeddings from the IMGT/mAbOnco-KG, we first flatten the KG format in tabular format with three columns corresponding to entities (*h* and *t*) and relations (*r*), then we preprocess it. The preprocessing includes shorting the URIs using namespaces (for example: *https://www.imgt.org/imgt-ontology#* will be replaced with the namespace imgt:) and cleaning the data by removing missing values, textual annotations, literals and keeping only the entities and their relation. Once the preprocessing is complete, we split the KG triples into different subsets using the ratios 0.8*/*0.1*/*0.1 with 80% for the training set, 10% for the validation set and 10% for the test set.

Table 1 provides statistics about IMGT/mAbOnco-KG and the splitting process.

**Table 1.** Statistics about IMGT/mAbOnco-KG

| Metric | Value |
| --- | --- |
| Triples | 29,795 |
| Entities | 9,611 |
| Relations | 53 |
| Training set size | 24,135 |
| Validation set size | 2,856 |
| Testing set size | 2,804 |

### 3.2 Link Prediction Problem Formulation in MAbs Repurposing

The task of mAb repurposing consists of proposing mAbs in the advanced clinical phase: phase M (approved) or phase III (pre-approval phase) in new clinical indications or new therapies. Thus, the approved mAb can be used in new pathologies. This is justified by different facts, including the following:

- The pathway overlap: different diseases can share common molecular mechanisms, signalling cascades, or immune dysregulation pathways, consequently, mAbs initially developed for one indication may be effective in another if they target a shared pathological process.
- The mechanism of action: a mAb’s target can be involved in different diseases, in this case, the mAb could be used in these diseases. For example, ranibizumab, an anti-VEGFA (vascular endothelial growth factor A) mAb initially developed for ophthalmology, is now used for age-related macular degeneration because of its role in the angiogenesis [36].

The mAb repurposing task can be formulated as a link prediction problem in which we generate scientific hypotheses on the reuse of existing mAbs in new clinical indications. For instance, in Figure 6, given two mAbs A and B, *can the mAb B, initially indicated in a clinical indication B, be used in a clinal indication A? or can the mAb A, initially indicated in a clinical indication A, be used in a clinal indication B?*. The way to answer this question is to predict the link between mAb B and its potential new clinical indications. Consequently, for a given mAb, we could predict a list of its potential clinical indications or for a given clinical indication, we could predict the potential list of associated mAbs.

**Fig. 6.**
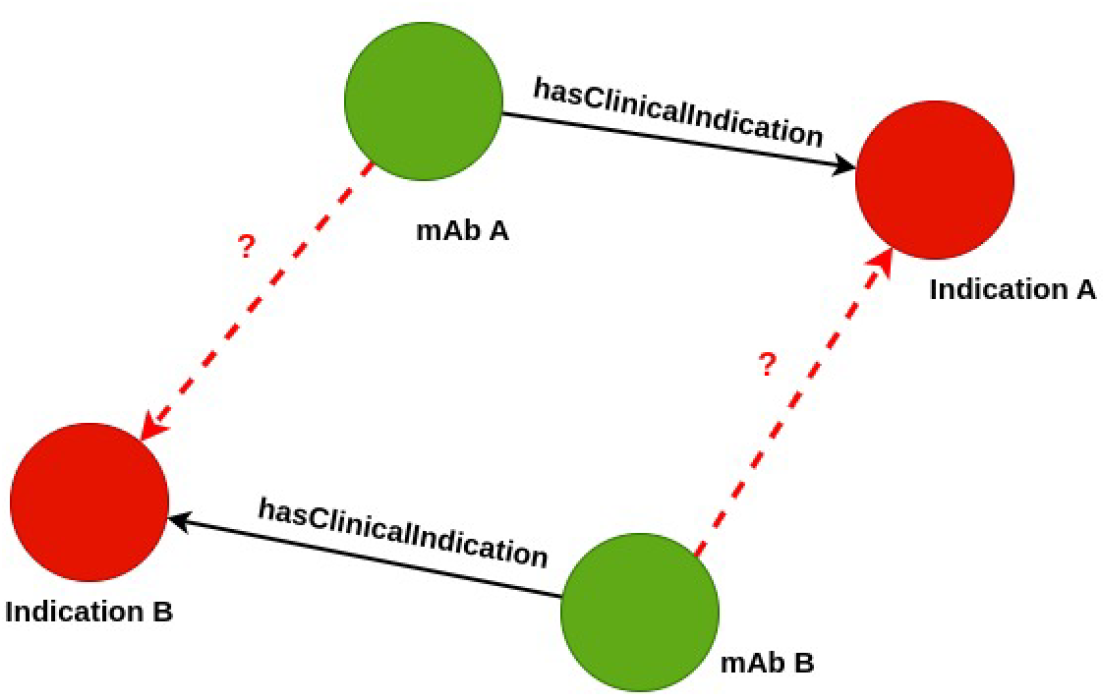
Formulation of the link prediction problem in the case of mAb repurposing

### 3.3 Evaluation of KGE models

To evaluate the generated embeddings in a link prediction setting, three common ranking metrics are used:

- The Mean Rank (MR) is used to evaluate a model’s ability to predict the correct position of entities or relations in triplets. It is computed as the average rank of correct predictions across the dataset, with lower values indicating better performance. While MR provides a general measure of prediction quality, it is sensitive to outliers, as incorrect predictions with high ranks can disproportionately affect the result. It is defined as follows:

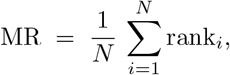

where rank_*i*_ is the rank of the true entity (head or tail) for the *i*-th test triplet among all possible candidates in the evaluation set N.
- The Mean Reciprocal Rank (MRR), evaluates the average of the reciprocals of the ranks of the first correct predictions for each triplet, emphasizing precision by rewarding models that rank correct answers higher. A higher MRR reflects a model’s ability to place correct predictions near the top, making it particularly useful for applications requiring precise initial predictions. However, MRR is limited because it considers only the first correct prediction and ignores others, which can skew its assessment of ranking quality. Its formula is:

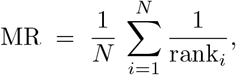
- The HITS@*k* metric evaluates the percentage of cases where a correct prediction appears within the top *k* predictions, providing a flexible measure of model performance. It is intuitive and straightforward, focusing on whether the correct answer is among the top *k* rather than its exact rank. While the HITS@*k* metric is tolerant and effective for assessing general ranking quality, it does not distinguish between the ranks within the top *k*, making it less precise than rank-sensitive metrics such as MRR. HITS@k is defined as follows:

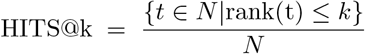

where {*t* ∈ *N* rank(*t*) ≤ *k*} is the subset of test triples whose correct entity is ranked within the top k.

Because we want to develop a system that prioritizes the ranking of the most likely mAb-clinical indication interactions (MRR) and identifies multiple plausible mAbs candidates (HITS@*k*), we define a new metric reflecting a weighted evaluation of the model’s overall performance called the performance score or *MRR*@10. This new metric combines the precision of the MRR metric and the coverage of HITS@*k*. It also simplifies model comparison. We chose HITS@10 for its ability to capture a broader ranking of predictions. The formula of the MRR@10 is described below, we set *α* to 0.6 or 60% for MRR and 0.4 or 40% for HITS@10.

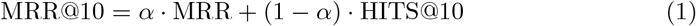

The *MRR*@10 metric aims to improve comparisons by offering a balanced and comprehensive measure of model performance, addressing limitations of individual metrics.

### 3.4 Experiments Settings

In our study, we chose the python PyKEEN tool [37, 38], which implements 40 embedding models, 15 loss functions and multiple reference datasets. With a very active community, the PyKEEN tool is flexible and easy to use, and is currently positioned as one of the references in the field of KG representation.

First, we implemented 28 KGE models from PyKEEN to select the top ten best models by considering a representation dimension or embedding size of 100. We used the models as they are provided in PyKEEN with their best default hyperparameters. They are trained on 500 epochs using a training batch of 64, and are validated every 100 epochs with the validation set. In this first experiment, we compare two strategies for generating the corrupted triplet set: random generation or basic generation and the Bernoulli strategy using probability for corrupted triplet generation. While the first strategy generates negative triplets randomly, the second generates negative triplets on the basis of the total number of *h* and *t* via the formula 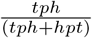 with *tph* the average number of *t* per *h* and *hpt* the average number of *h* per *t* [38]. For this experiment, we generate one negative triplet per positive triplet.

In this first experiment, we selected the top ten models using the most effective corruption strategies. We then tested these models with different embedding sizes and varying numbers of negative triplets generated. Specifically, we evaluated our ten best models across four embedding dimensions (100, 200, 400, and 800) and four settings for the number of negative triplets generated (1, 64, 128, and 256). Table 2 presents the different combinations of embedding sizes and the number of negative triplets generated. For example, we tested embedding sizes ranging from 100 to 800 with one negative triplet generated for each positive triplet. The aim was to identify the optimal settings for each model by considering both the embedding dimensions and the negative sampling parameters.

**Table 2.** Combinations of the number of negative triplets and embedding sizes.

| Neg/Pos Triplets | Embedding Sizes |
| --- | --- |
| 1 | (100, 200, 400, 800) |
| 64 | (100, 200, 400, 800) |
| 128 | (100, 200, 400, 800) |
| 256 | (100, 200, 400, 800) |

To develop a scientific hypothesis generation system, that generates mAb candidates for a given clinical indication, we consequently evaluated the best KGE models on this task by using their best settings previously found. For that, in our testing and validation set, we consider only relations involving clinical indications and mAbs, that is clinical indication relations (*imgt:hasClinicalIndication* and its reverse *imgt:isClincicalIndicationOf*). This restricted evaluation will allow us to determine how well a KGE model fits our particular task and will guide the choice of the model for the user.

## 4 Results

### 4.1 Experiment Results

The first experiment, consisting of the evaluation of two strategies of triplet corruption, including basic, and Bernoulli strategies, allows us to select the top ten best models with the best strategy from the 28 models evaluated. To compare these strategies, we measure the mean and max values of the different metrics: MRR, HITS@*k* and MRR@10, as shown in Figure 7. The analysis of the mean shows that the best strategy for generating the corrupt triplets is the Bernoulli strategy. In fact, on average, the Bernoulli strategy, yields the best negative triplets whether, it is the MRR (39.08 *>*37.19 for example) or the HITS@*k* (51.85*>*49.09 for the HITS@10). This analysis is also valid for the max measure of different metrics: the best strategy giving the best result is Bernoulli, for example, for the MRR@10 metric, we have 79.24*>*76.39 or 82.94 *>* 80.03 for the HITS@10.

**Fig. 7.**
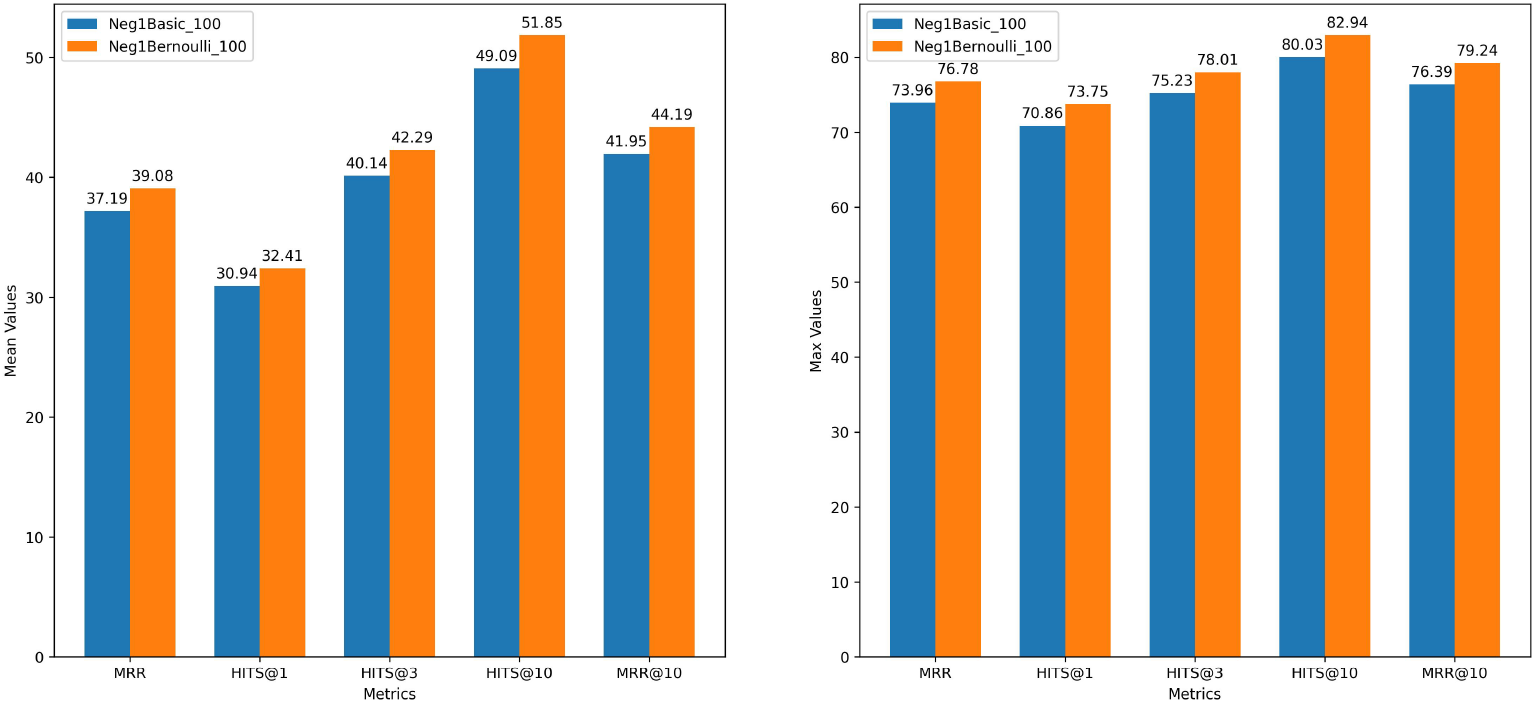
Comparison of the two strategies of triplets corruption

Figure 8 shows the evaluation of the 28 models using our new metric MRR@10 in the Bernoulli corruption strategy. The top ten best models are: *PairRE, TorusE, RotatE, DistMult, ComplEx, BoxE, TransE, TransD, CrossE, and SimplE*. These best models are selected for further experiments.

**Fig. 8.**
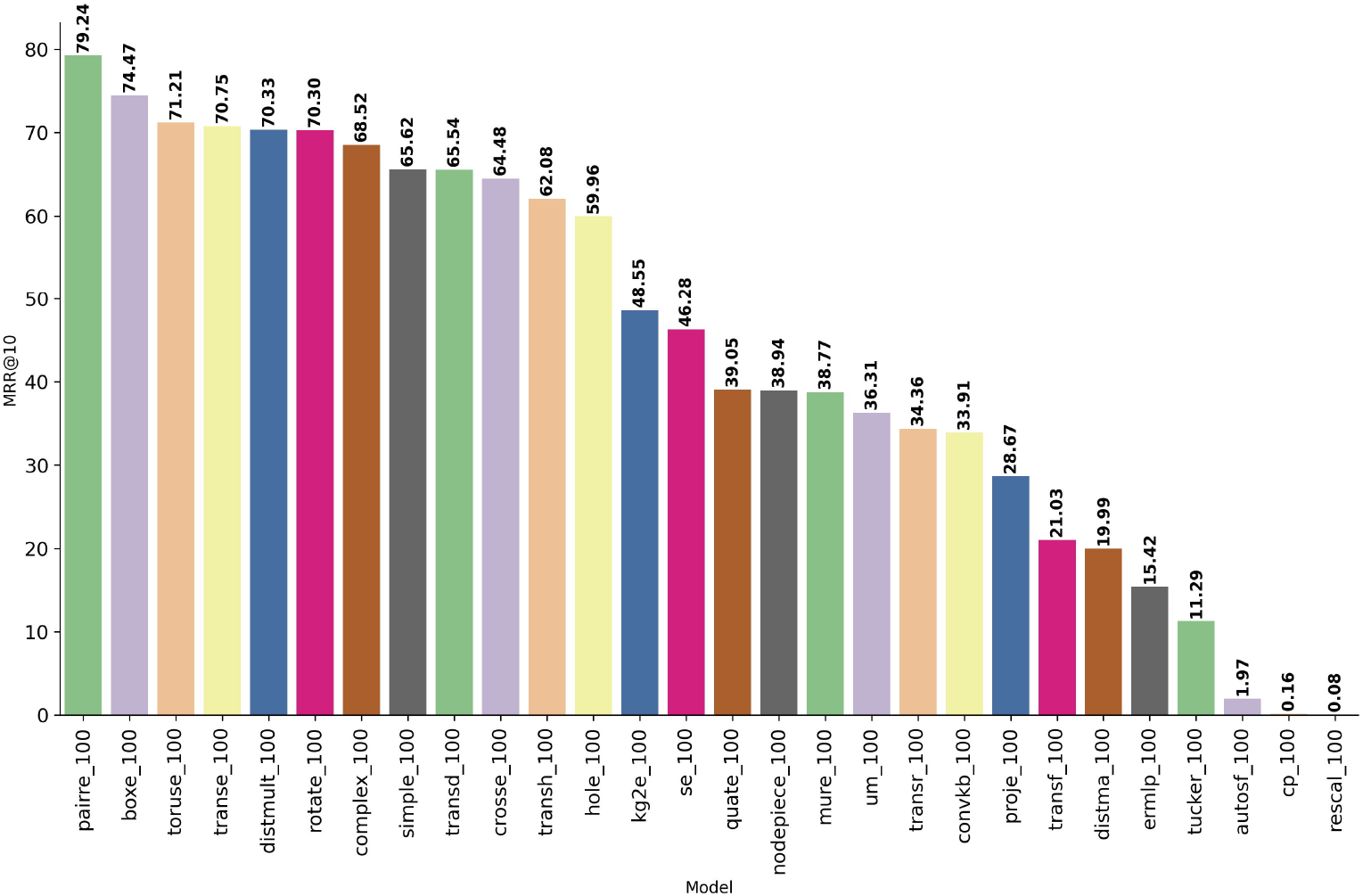
Evaluation of the 28 models with the Bernoulli corruption strategy using the MRR@10 metric

In the second experiment, we combined the number of embedding sizes with the number of negative triplets generated by positive triplet via the Bernoulli strategy (Table 2). Table 3 provides the best settings for each model, the top three model based on the MRR@10 metric (Figure 9) are:

**Table 3.**
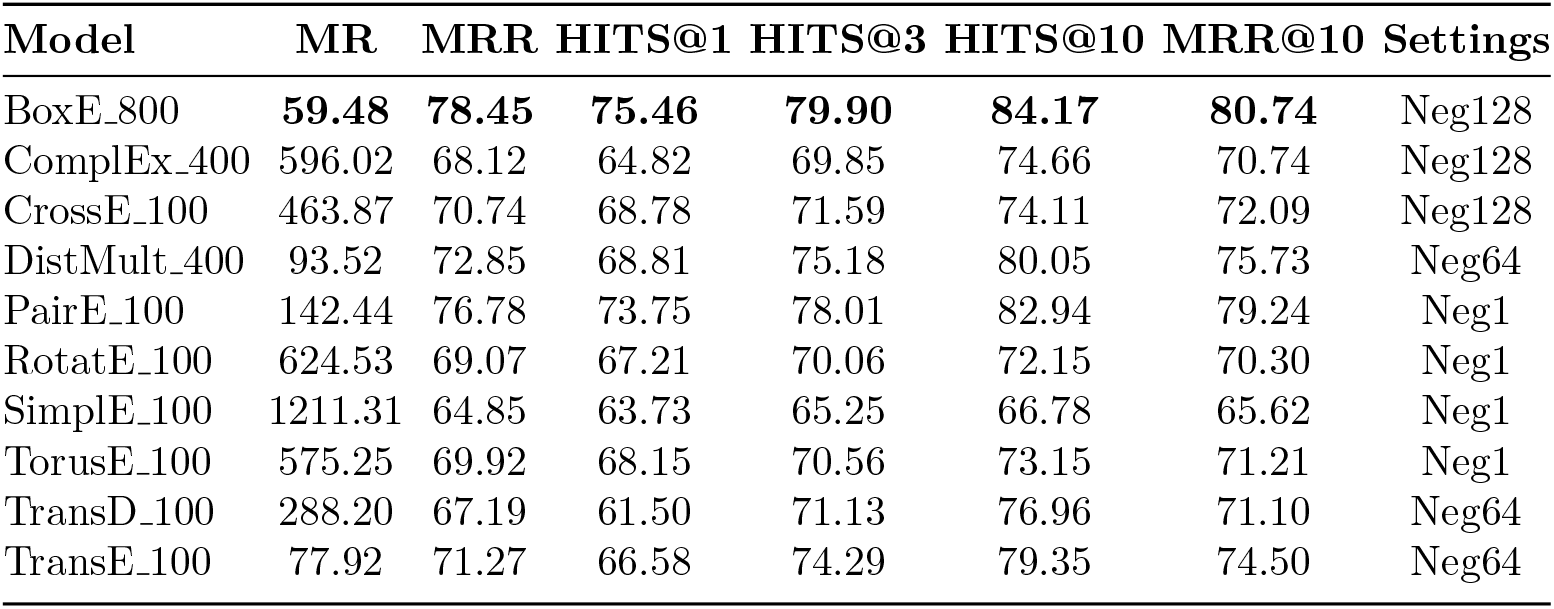
Best Settings for each models

| Model | MR | MRR | HITS@1 | HITS@3 | HITS@10 | MRR@10 | Settings |
| --- | --- | --- | --- | --- | --- | --- | --- |
| BoxE_800 | **59.48** | **78.45** | **75.46** | **79.90** | **84.17** | **80.74** | Neg128 |
| ComplEx_400 | 596.02 | 68.12 | 64.82 | 69.85 | 74.66 | 70.74 | Neg128 |
| CrossE_100 | 463.87 | 70.74 | 68.78 | 71.59 | 74.11 | 72.09 | Neg128 |
| DistMult_400 | 93.52 | 72.85 | 68.81 | 75.18 | 80.05 | 75.73 | Neg64 |
| PairE_100 | 142.44 | 76.78 | 73.75 | 78.01 | 82.94 | 79.24 | Neg1 |
| RotatE_100 | 624.53 | 69.07 | 67.21 | 70.06 | 72.15 | 70.30 | Neg1 |
| SimpleE_100 | 1211.31 | 64.85 | 63.73 | 65.25 | 66.78 | 65.62 | Neg1 |
| TorusE_100 | 575.25 | 69.92 | 68.15 | 70.56 | 73.15 | 71.21 | Neg1 |
| TransD_100 | 288.20 | 67.19 | 61.50 | 71.13 | 76.96 | 71.10 | Neg64 |
| TransE_100 | 77.92 | 71.27 | 66.58 | 74.29 | 79.35 | 74.50 | Neg64 |

**Fig. 9.**
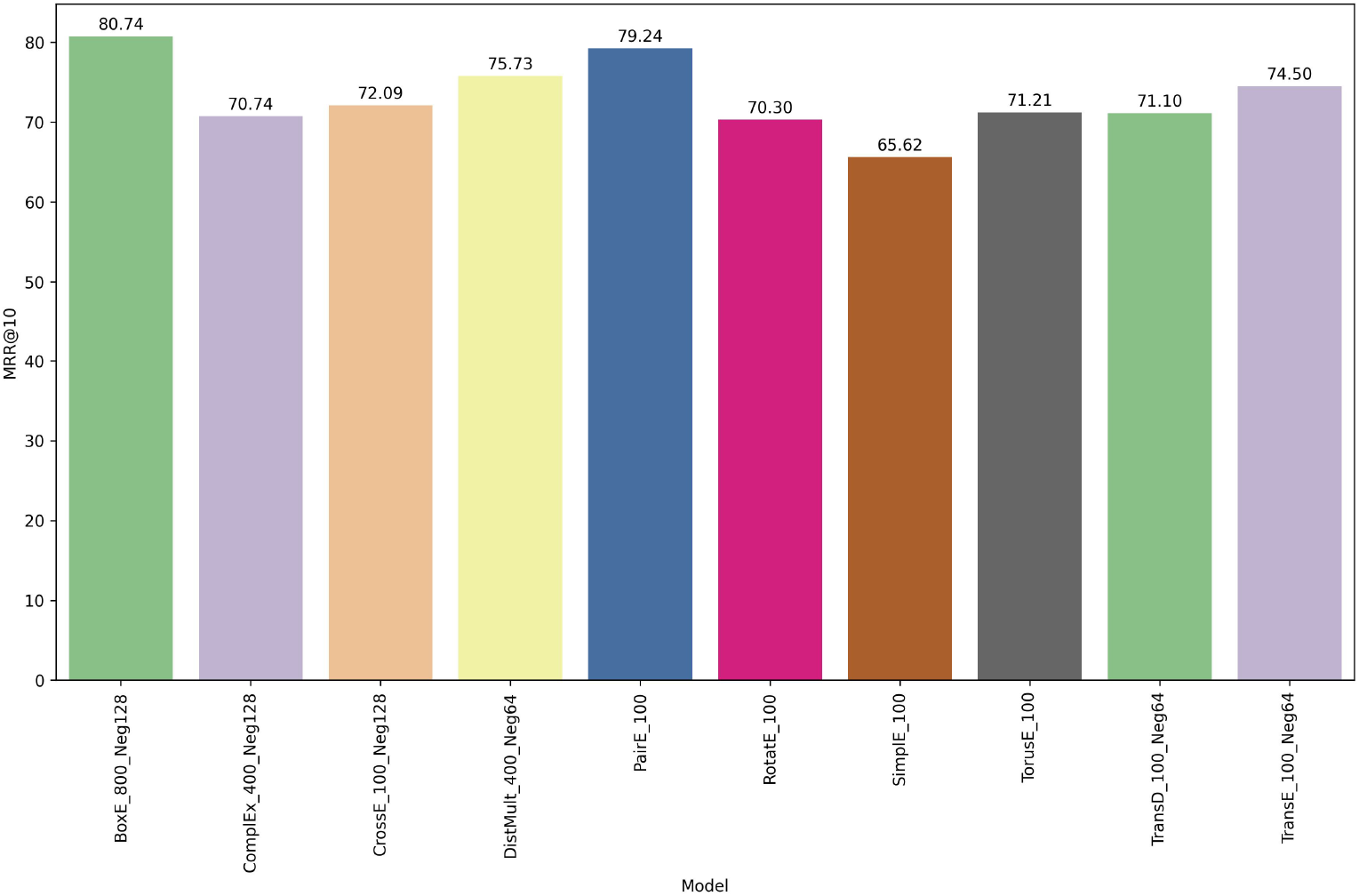
Best Settings for each model based on the MRR@10 metrics

- *BoxE* with the embedding dimension 800 and 128 negatives per positive triplet
- *PairRE* with the embedding dimension 100 and one negative per positive triplet
- *DistMult* with the embedding dimension 400 and 64 negatives per positive triplet

This analysis is valid for the metrics MRR, HIT@3 and the HIT@10 and the best three models based on the MR metrics are *BoxE, TransE* and *DistMult*.

For the final experiment, we assessed the top ten models on a restricted subset of the testing and validation data, focusing on relations involving mAbs and clinical indications. This subset accounts for 12.95% of the overall testing and validation dataset.

Table 4 shows the result of this evaluation, and the *BoxE* model again yields the best results for the MR, MRR, MRR@10 and HITS@*k* metrics.

**Table 4.**
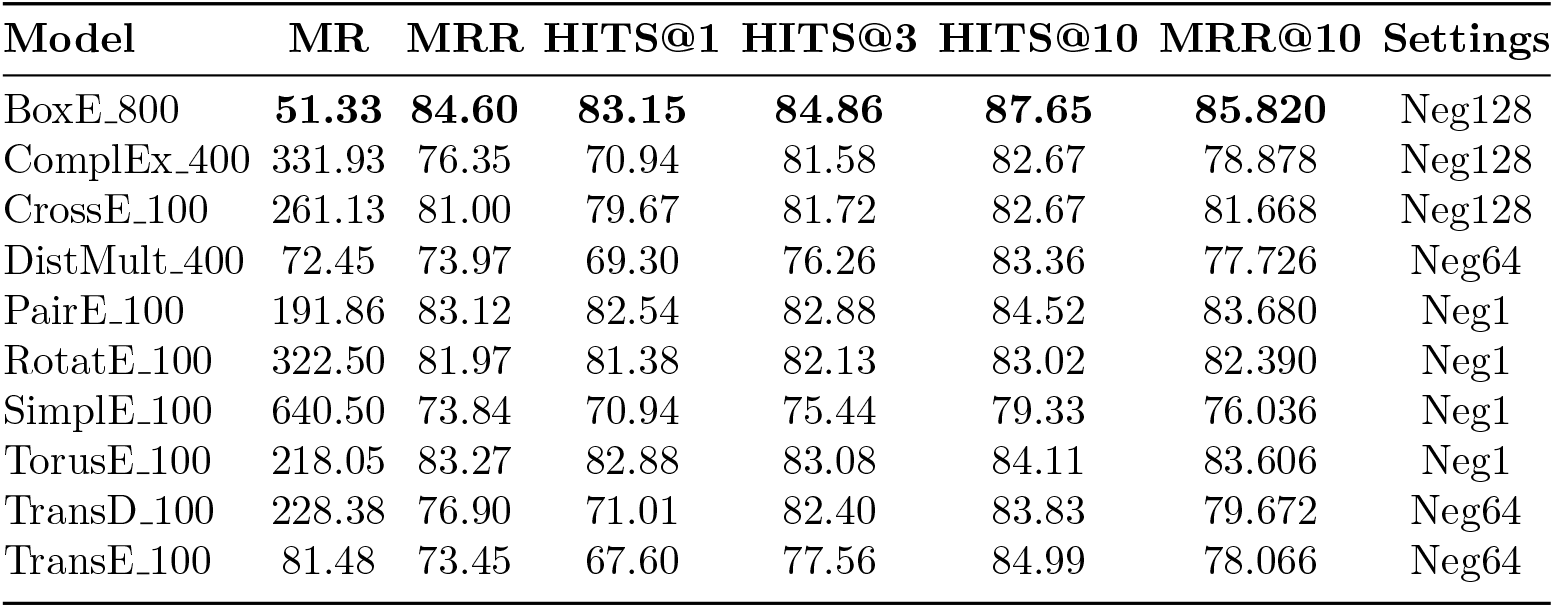
Evaluation of the best settings of the models in a data restriction configuration, considering the clinical indications

These experiments indicate that regardless of the metrics, the *BoxE* model performs better than the other nine models do. This superior performance of *BoxE* is due to its representation approach which uses boxes to capture the semantic of the KG. In fact, *BoxE*, represents entities and relations using hyper-rectangles or boxes in a high-dimensional space, allowing it to effectively capture asymmetry, hierarchy, and relational composition [39]. This structured representation enables *BoxE* to model complex relation types, including one-to-many and many-to-many relationships, with greater flexibility and accuracy. Additionally, *BoxE* efficiently encodes relational constraints, making it particularly effective in link prediction tasks.

### 4.2 Visualisation of the embeddings

Since human sensory perception and conventional graphical representations are typically confined to three dimensions or fewer, visualizing high-dimensional embeddings poses a challenge. To overcome this and effectively assess the quality of our generated embeddings, we applied dimensionality reduction techniques, such as t-distributed stochastic neighbor embedding (t-SNE)[40], to project them into two and three dimensions for visualization. We then plotted the reduced embeddings to evaluate the model’s ability to capture and distinguish KG entities effectively. For example, Figure 10 presents the 2D visualizations of the embeddings generated by the best KGE model:*BoxE*. This figure shows that the different categories are well separated, indicating the best quality of the embeddings generated by the *BoxE* model on our KG. In fact, we can clearly see the different clusters of the mAbs (green), targets (cyan), clinical indications (red) or study products (soft pink). Some entities belong to the construct (green light) and segment (brown) clusters because a construct is made of at least one segment, consequently the construct can be made of only one segment 10.

**Fig. 10.**
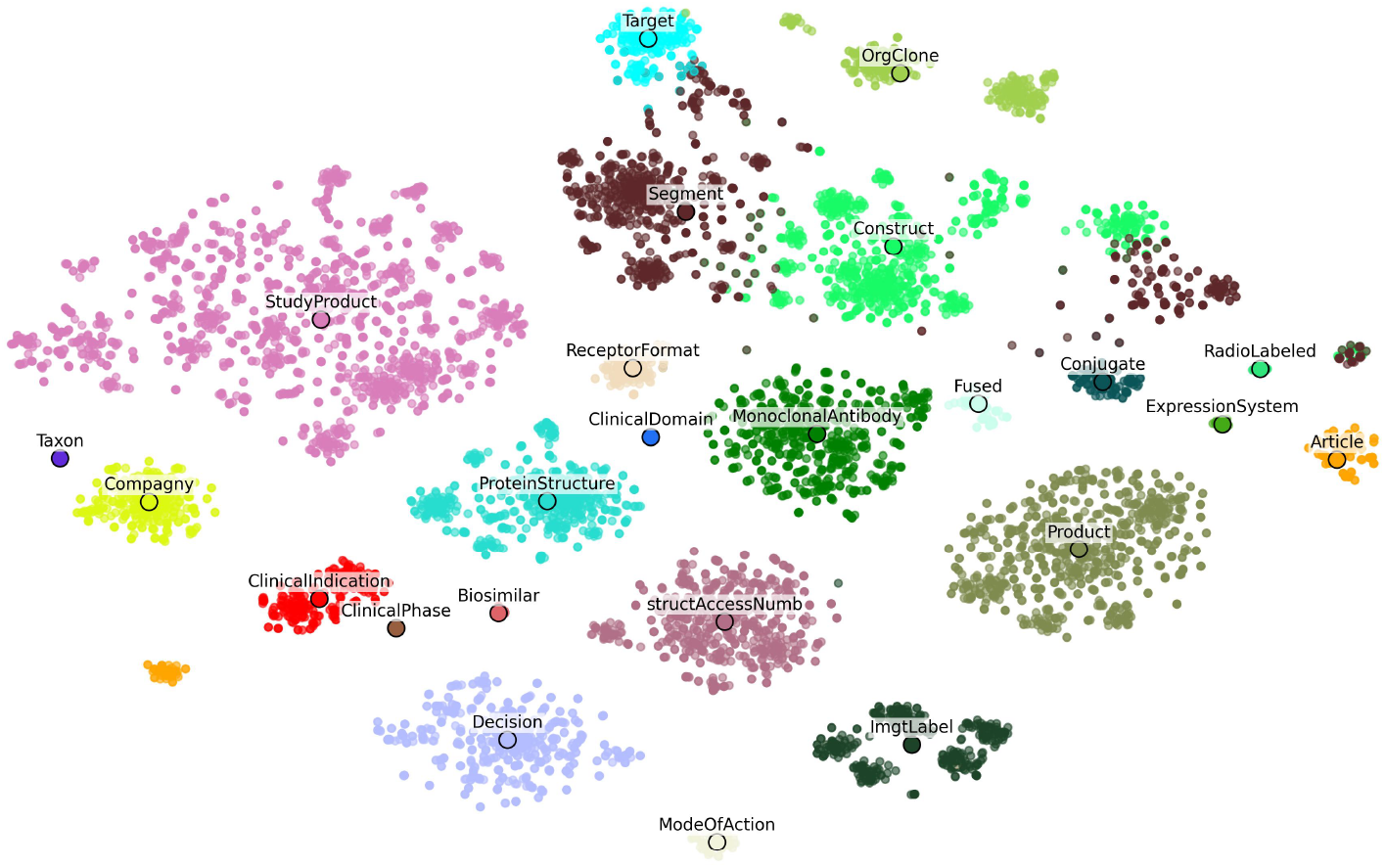
2D t-SNE visualization of *BoxE* embeddings with d = 800 and 128 negative triplets per positive

### 4.3 Application of hypothesis generation in mAbs repurposing

With these ten models and their best settings, we develop an application for generating scientific hypotheses in mAb repurposing. Our application takes as input a clinical indication in the oncology domain, and transfers it to a selected KGE model, which will generate a list of potential mAb candidates susceptible to be used in the selected clinical indication (Figure 11).

**Fig. 11.**
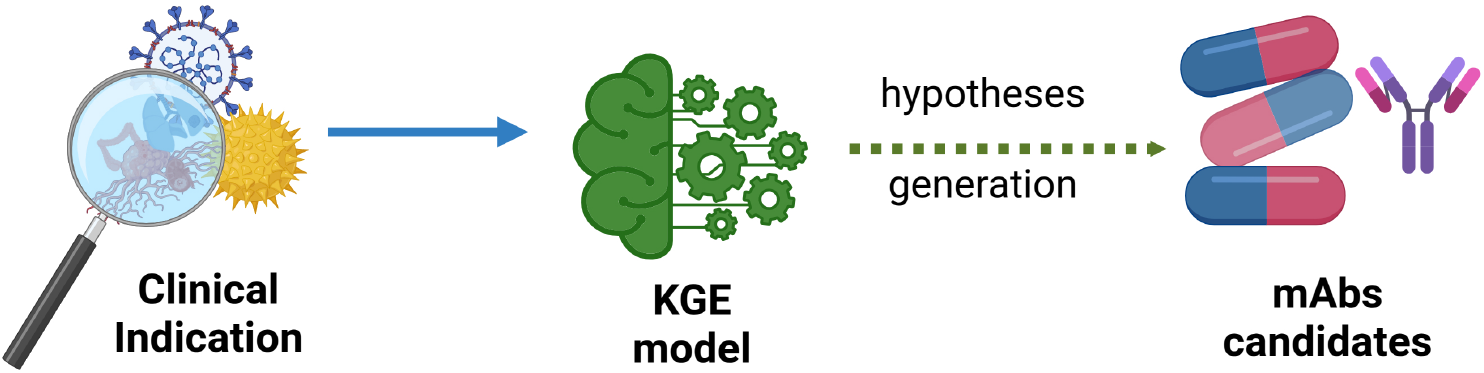
Generating mAbs candidates for a given target or clinical indication

The developed application is integrated into our latest mAb exploration application accessible at https://imgt.org/mAb-KG/. A new interface (https://imgt.org/mAb-KG/Hypotheses_Generation) dedicated to the mAb candidate generation, provides access to the ten previous KGE models and embeddings visualizations. It allows selecting the clinical indication for the mAbs candidate generation. On the interface, a list of clinical indication entities is proposed, and a clinical indication can be selected to generate scientific hypotheses. The interface allows the generation of one to 20 mAb candidates for a given clinical indication, and the user can filter the generation by removing the information that appeared in the training. For example, Figure 12 provides an example of the hypotheses generated in the case of the prostate cancer via the *BoxE* model. All the suggested mAbs are already used in the prostate cancer (Figure 13). The score value depends on the model and cannot be interpreted as a probability and cannot be used for model comparison.

**Fig. 12.**
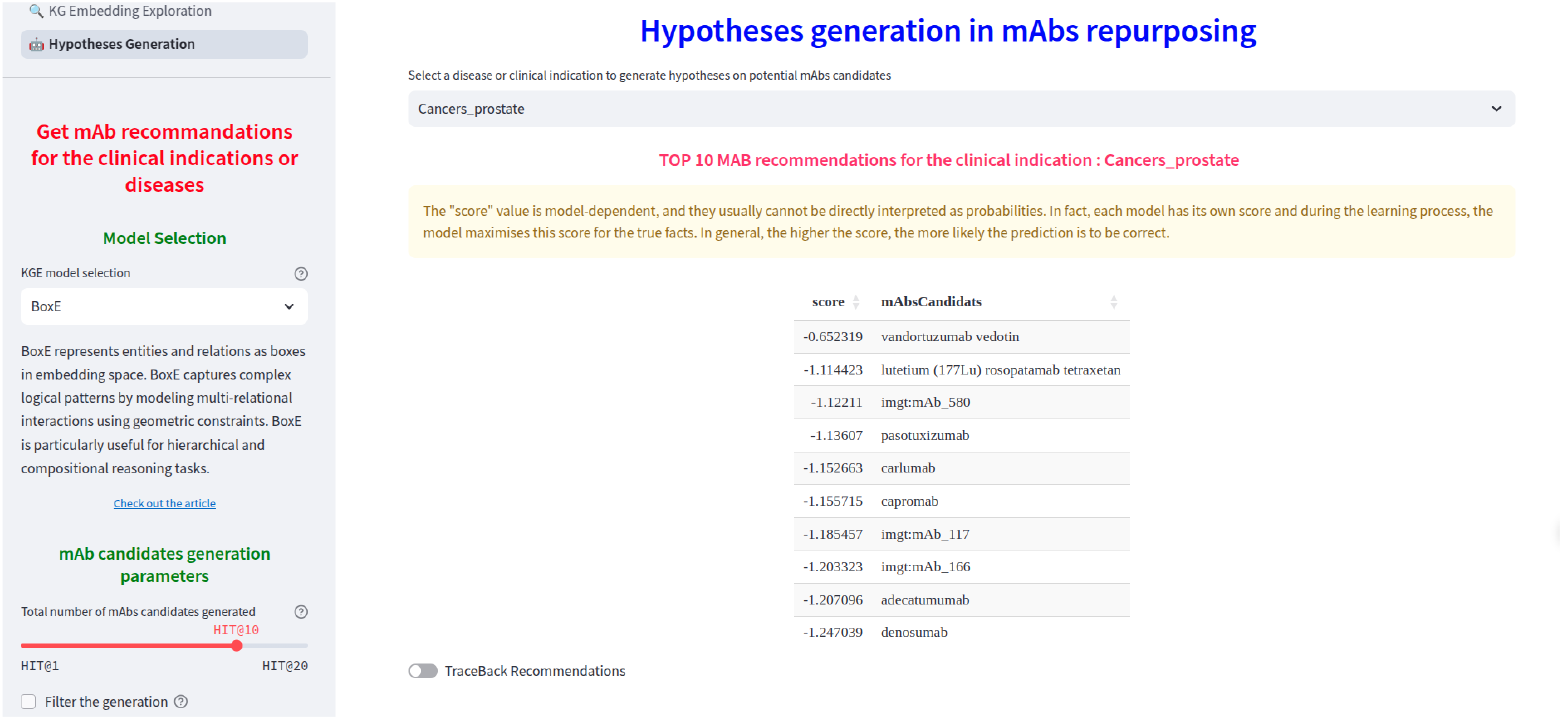
Hypotheses generation for the prostate’s cancer: top ten mAbs candidates

**Fig. 13.**
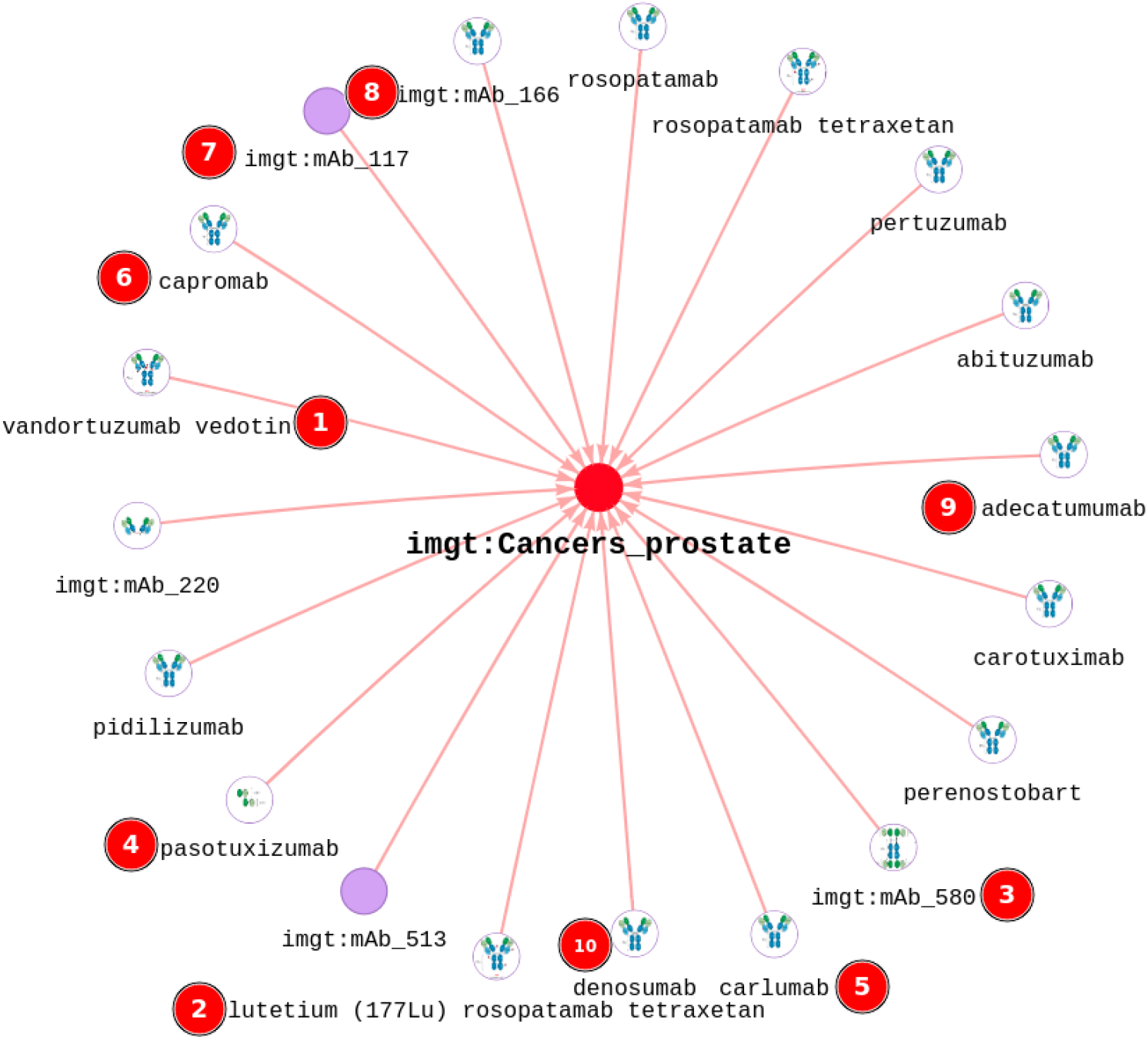
Real mAbs used to treat prostate’s cancer: in red circle with number, the candidates generated by the *BoxE* model

Given the explainability support that a KG provides to the machine learning prediction system [41], we trackback the hypotheses generated (mAb candidate generation) by the models (Figure 14). This tracing consists of taking the list of mAb candidates generated and retrieving them in real time via queries in IMGT-KG. Then, we display the graphs associated with the queries, allowing us to visualize the generated candidates and their different connections. This traceback can serve as a visual and interpreting support for the mAbs candidates generated. Figure 14 shows details of the predicted mAbs, as in [12], the user can add additional levels including the construct and the mode of action of the mAbs generated. These details provide a complete support to interpret the hypotheses generated.

**Fig. 14.**
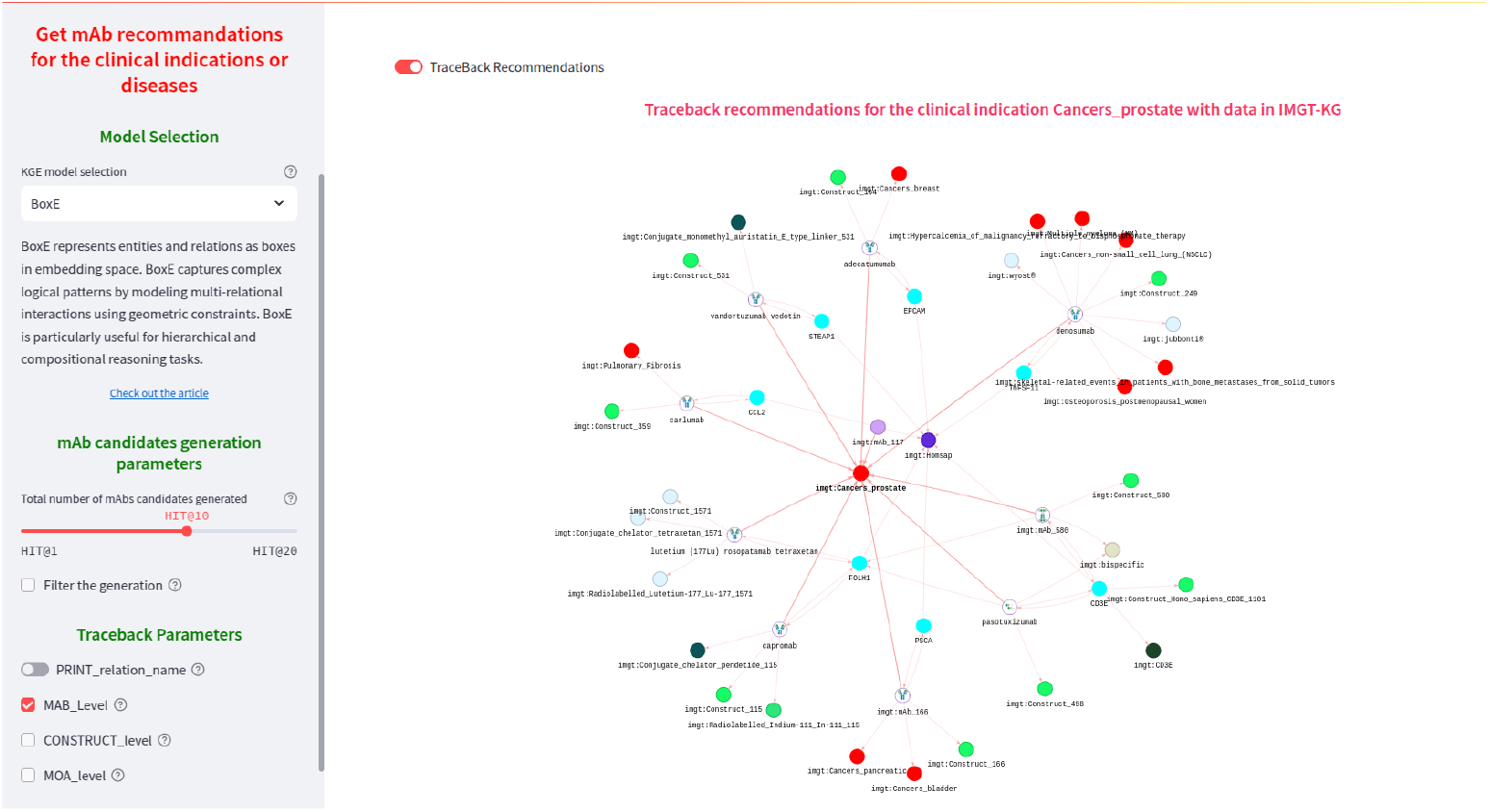
Visual support of the mAb candidates predicted to treat the prostate’s cancer

We also provide an additional interface for exploring the embeddings generated by the KGE models at https://imgt.org/mAb-KG/KG_Embedding_Exploration. This interface provides a two-dimensional and three-dimensional visualisation tool for the generated embeddings based on four dimensional reduction algorithms including t-SNE, PCA [42], Trimap [43] and UMAP.

## 5 Selected Scientific Hypotheses Generated in mAbs Repurposing

We used our tool to generate scientific hypotheses for the repurposing of existing mAbs for two clinical indications: lymphocytic leukemia (CLL) and ovarian cancer.

### 5.1 Ovarian cancers

Ovarian cancer is the seventh most commonly diagnosed cancer among women in the world [44]. It is a global health problem diagnosed at a late stage and has no effective screening strategy [45]. In fact, ovarian cancer remains the most lethal gynecological cancer, despite the recent advances in its treatment [46]. Aiming to improve the patient outcome, various mAbs (avelumab, ubamatamab, olaratumab, etc) with promised results, have been used to treat the ovarian cancer [46]. Consequently, generating hypotheses to repurpose existing mAbs to treat this cancer should accelerate the management of the disease.

We subsequently used our tool, to generate mAb candidates susceptible to be used in ovarian cancer. We used the *BoxE* model to generate hypotheses on mAbs repurposing by considering the clinical indication *Cancers ovarian*. We filter the generated mAbs list by removing known triplets that appear in the training set. As a result, we obtained the following top ten predicted mAbs: lifastuzumab vedotin, abituzumab, avelumab, catumaxomab, imgt:mAb 736, imgt:mAb 98, aflibercept, nimotuzumab, atezolizumab, olaratumab. While eight suggested candidates are already known to be used in the treatment of the ovarian cancer, the two other predicted such as atezolizumab, and olaratumab are not used in to treat ovarian cancer. In the case of atezolizumab, the prediction traceback shows that it targets the CD274 (PD-L1) protein which is also a target of avelumab (initially proposed to treat ovarian cancer).

Indeed, atezolizumab, a mAb targeting CD274 (PD-L1), has been investigated for the treatment of ovarian cancer, particularly in combination with chemotherapy or bevazucimab (anti-VEGF). Given the significant role of immune evasion in ovarian cancer progression, clinical trials have evaluated the potential of atezolizumab to improve patient outcomes, especially in advanced or recurrent cases [47–49]. While these studies have shown that adding atezolizumab to standard chemotherapy and bevacizumab does not provide a significant clinical benefit in ovarian cancer, it would be interesting to explore its effects without combination since it targets CD274 which is also targeted by avelumab initialy used to treat ovarian cancer.

### 5.2 Scientific Hypotheses Generated in Chronic Lymphocytic Leukemia

Chronic lymphocytic leukemia (CLL) is the most common form of leukemia in Western countries [50, 51]. It is characterized by clonal proliferation and accumulation of neoplastic B lymphocytes in the blood, bone marrow, lymph nodes, and spleen [50]. Owing to the palliative nature of chemotherapy, the use of mAbs in the treatment of CLL has extensively increased [51]. In fact, various mAbs including rituximab, alemtuzumab or ofatumumab have been used as therapeutic option for CLL [51]. In CLL disease, the most targeted proteins are MS4A1 or CD20 (24 studies) and CD19 (16 studies) (Figure 5) expressed on B-cells. Generating hypotheses concerning the repurposing of existing mAbs as a therapeutic option could enhance the management and treatment of CLL.

We used the *BoxE* model to generate potential mAb candidates that can be used in CLL therapy. We explored the top 15 generated candidates: apolizumab, inebilizumab, polatuzumab vedotin, brexucabtagene autoleucel, lucatumumab, rituximab, alemtuzumab, zeripatamig, ublituximab, ofatumumab, zilovertamab, odronextamab, glofitamab, interferon alfa and loncastuximab tesirine. The first thirteen candidates are already known for use in the CLL therapy, whereas the last three (glofitamab, interferon alfa and loncastuximab tesirine) are not known for use in the treatment of CLL.

Both loncastuximab tesirine (anti-CD19) and glofitamab (anti-MSA41 or anti-CD20 and anti-CD3E) are mAbs used to treat B-cell malignancies, including certain types of non-Hodgkin lymphoma (NHL). Although, their use in CLL is less established, they have the same targets (CD19, MS4A1 and CD3E) as mAbs designed for CLL such as rituximab, odronextamab, ofatumumab or inebilizumab.

Glofitamab is a bispecific mAb that simultaneously targets CD20, including CLL cells and CD3 expressed on T-cells, leading to T-cells activation and proliferation [52]. Consequently, the T-cell will engage and kill CD20-positive B-cells. Since CD20 is expressed in the CLL, glofitamab’s mechanism of action could theoretically be effective in CLL.

Loncastuximab tesirine is an antibody-drug conjugate with pyrrolobenzodiazepines (PBD) targeting the CD19 [53]. The protein CD19 plays a critical role in B-cell receptor signalling and is also expressed on the surface of CLL cells. When loncastuximab tesirin binds to its target on CD19-positive cells, it is internalized and then delivers the PBD inside the cell, leading to cell death [53]. Considering that CD19 is expressed in CLL, the mechanism of action of loncastuximab tesirine suggests potential applicability in CD19-positive CLL cases.

Moreover, both loncastuximab tesirine and glofitamab are currently under investigation in clinical trials for CLL conducted by independent research groups. According to publicly available data, glofitamab is being evaluated in a [phase II trial](https://clinicaltrials.gov/study/NCT06043674), and loncastuximab tesirine is being evaluated in a [phase I trial](https://clinicaltrials.gov/study/NCT05971251).

## 6 Conclusion

Over the past few years, mAbs have revolutionized the field of targeted cancer therapy. They constitute a promising class of targeted anticancer agents that enhance natural immune system functions to suppress cancer cell activity, and eradicate cancer cells [54]. Given the interest in mAbs and their vast medical capabilities, we developed IMGT/mAb-KG, the IMGT-KG for therapeutic monoclonal antibodies [12], which provides descriptions about 1,586 mAbs and their related knowledge including targets, clinical indications, mode of actions etc. Due to the relatively recent nature of mAbs in comparison with current treatments, a considerable amount of research remains to be conducted and discoveries to be made [3]. Furthermore, the current wet lab-based development of drugs or mAbs is time-consuming, labor-intensive, and costly, with a low success rate. A parallel investigation, which is less time-consuming, is exploring the repurposing of existing mAbs for new therapeutic applications.

To accelerate the drug repurposing process, we propose a scientific hypothesis tool for generating potential mAb candidates by using an oncology dedicated subset of IMGT/mAb-KG available at https://imgt.org/mAb-KG/. This application generates potential mAb candidates susceptible to be used in a particular clinical indication. However, since the generation process is based on probabilities, its validity must be confirmed experimentally by biologists specialized in the field of mAb engineering. The generation process uses ten KGE models trained on the oncology subset of IMGT/mAb-KG with the best performances. In addition to the prediction of the mAbs candidate generation, we also provide a trackback system aiming to serve as visual support on the generated mAb candidates, thus providing a guide to the interpretability of the hypotheses. In fact, this visual support proposes a graph of the mAb candidates and their interactions in the IMGT/mAb-KG knowledge graph, allowing the user to compare the prediction with the real data. As use cases, with our tool, we generated scientific hypotheses for the treatment of ovarian cancers and CLL. In the case of CLL, we identify two mAb candidates such as loncastuximab tesirine and glofitamab. These two mAb candidates are currently in clinical trials for the treatment of CLL.

Our mAb candidate generation system currently functions in a transductive setting (fixed length of entities and relations in the KG), and thus, cannot be used for unseen entities (out of the KG). In our future works, we will explore foundational models including Ultra [55] or KG-ICL [56] to face this limitation by building an inductive mAbs candidates generation system and embedding all the IMGT-KG knowledge, then providing a KGE foundational model in immunogenetics.

## Declarations

### Ethics approval and consent to participate

Not applicable

### Consent for publication

All authors approved the manuscript and gave their consent for submission and publication.

### Availability of data and material

https://imgt.org/mAb-KG/Hypotheses_Generation and https://src.koda.cnrs.fr/imgt-igh/oncomabkgembeddings

## Acknowledgements

This project was provided with computing AI and storage resources by GENCI at IDRIS thanks to the grant 2024-AD011014321 on the super-computer Jean Zay’s A100 partition. We acknowledge the support of Immun4Cure University Hospital Institute “Institute for innovative immunotherapies in autoimmune diseases” (France 2030 /ANR-23-IAHU-009). We thank all members of the IMGT® team for their expertise and constant motivation. IMGT® is member of the French Infrastructure Institut Français de Bioinformatique (IFB) as well as member of BioCampus, MAbImprove and IBiSA.

## Clinical trial number

Not applicable.

## References

[1] Bray, F. et al. Global cancer statistics 2022: GLOBOCAN estimates of incidence and mortality worldwide for 36 cancers in 185 countries. CA: A Cancer Journal for Clinicians 74, 229–263 (2024).

[2] Sung, H. et al. Global Cancer Statistics 2020: GLOBOCAN Estimates of Incidence and Mortality Worldwide for 36 Cancers in 185 Countries. CA: A Cancer Journal for Clinicians 71, 209–249 (2021). URL https://onlinelibrary.wiley.com/doi/full/10.3322/caac.21660 https://onlinelibrary.wiley.com/doi/abs/10.3322/caac.21660 https://acsjournals.onlinelibrary.wiley.com/doi/10.3322/caac.21660.

[3] Srirapu, S. Monoclonal Antibodies and their Applications in Cancer. Journal of Student Research 12, 1–7 (2023).

[4] Castelli, M. S., McGonigle, P. & Hornby, P. J. The pharmacology and therapeutic applications of monoclonal antibodies. Pharmacology research & perspectives 7, e00535 (2019).

[5] Dos Santos, M. L., Quintilio, W., Manieri, T. M., Tsuruta, L. R. & Moro, A. M. Advances and challenges in therapeutic monoclonal antibodies drug development (2018).

[6] Ecker, D. M., Jones, S. D. & Levine, H. L. The therapeutic monoclonal antibody market (2015). URL http://dx.doi.org/10.4161/19420862.2015.989042.

[7] Lu, R. M. et al. Development of therapeutic antibodies for the treatment of diseases. Journal of Biomedical Science 2020 27:1 27, 1–30 (2020). URL https://jbiomedsci.biomedcentral.com/articles/10.1186/s12929-019-0592-z.

[8] Golbeck, J. et al. The National Cancer Institute’s Thesaurus and Ontology. SSRN Electronic Journal (2003). URL https://papers.ssrn.com/abstract=3199007 https://www.ssrn.com/abstract=3199007.

[9] Raybould, M. I. et al. Thera-SAbDab: The Therapeutic Structural Antibody Database. Nucleic Acids Research 48, D383–D388 (2020).

[10] Abanades, B. et al. The Patent and Literature Antibody Database (PLAbDab): an evolving reference set of functionally diverse, literature-annotated antibody sequences and structures. bioRxiv 2023.07.15.549143 (2023). URL https://www.biorxiv.org/content/10.1101/2023.07.15.549143v1 https://www.biorxiv.org/content/10.1101/2023.07.15.549143v1.abstract.

[11] C., P., andGinestoux C., W. Y., Ehrenmann, D. P. & M.-P., L. IMGT/mAb-DB: the IMGT® database for therapeutic monoclonal antibodies (2010). URL http://www.imgt.org.

[12] Sanou, G. et al. IMGT/mAb-KG: the knowledge graph for therapeutic monoclonal antibodies. Frontiers in Immunology 15, 1393839 (2024). URL https://www.frontiersin.org/articles/10.3389/fimmu.2024.1393839/full.

[13] Mohamed, S. K., Nounu, A. & Novàček, V. Drug target discovery using knowledge graph embeddings. Proceedings of the ACM Symposium on Applied Computing Part F1477, 11–18 (2019).

[14] Park, K. A review of computational drug repurposing. Translational and Clinical Pharmacology 27, 59–63 (2019). URL 10.12793/tcp.2019.27.2.59.

[15] MacLean, F. Knowledge graphs and their applications in drug discovery. Expert Opinion on Drug Discovery 16, 1057–1069 (2021). URL 10.1080/17460441.2021.1910673.

[16] Boudin, M., Diallo, G., Drancé, M. & Mougin, F. The OREGANO knowledge graph for computational drug repurposing. Scientific Data 10, 1–13 (2023). URL https://www.nature.com/articles/s41597-023-02757-0.

[17] Mullard, A. New drugs cost US$2.6 billion to develop. Nature Publishing Group (2014).

[18] Ghorbanali, Z., Zare-Mirakabad, F., Akbari, M., Salehi, N. & Masoudi-Nejad, A. DrugRep-KG: Toward Learning a Unified Latent Space for Drug Repurposing Using Knowledge Graphs. Journal of Chemical Information and Modeling (2023).

[19] Cummings, S. R. et al. Denosumab for prevention of fractures in postmenopausal women with osteoporosis. The New England journal of medicine 361, 756–765 (2009). URL https://pubmed.ncbi.nlm.nih.gov/19671655/.

[20] Bhulani, N. et al. A phase 3 study to determine the breast cancer risk reducing effect of denosumab in women carrying a germline BRCA1 mutation (BRCA-P Study). Journal of Clinical Oncology 40, TPS10616–TPS10616 (2022). URL https://ascopubs.org/doi/10.1200/JCO.2022.40.16_suppl.TPS10616.

[21] Gimeno, A. et al. The Light and Dark Sides of Virtual Screening: What Is There to Know? International Journal of Molecular Sciences 20 (2019). URL /pmc/articles/PMC6470506//pmc/articles/PMC6470506/?report=abstracthttps://www.ncbi.nlm.nih.gov/pmc/articles/PMC6470506/.

[22] Adeshina, Y. O., Deeds, E. J. & Karanicolas, J. Machine learning classification can reduce false positives in structure-based virtual screening. Proceedings of the National Academy of Sciences of the United States of America 117, 18477–18488 (2020). URL https://www.pnas.org/doi/abs/10.1073/pnas.2000585117.

[23] Zhang, R. et al. Drug repurposing for COVID-19 via knowledge graph completion. Journal of Biomedical Informatics 115 (2021). URL http://arxiv.org/abs/2010.09600 http://dx.doi.org/10.1016/j.jbi.2021.103696.

[24] Islam, K. et al. Molecular-evaluated and explainable drug repurposing for COVID-19 using ensemble knowledge graph embedding. Scientific Reports — 13, 3643 (123). URL 10.1038/s41598-023-30095-z.

[25] Grohe, M. Word2vec, node2vec, graph2vec, X2vec: Towards a Theory of Vector Embeddings of Structured Data. Proceedings of the ACM SIGACT-SIGMOD-SIGART Symposium on Principles of Database Systems 1–16 (2020).

[26] Blake, J. A. & Bult, C. J. Beyond the data deluge: Data integration and bioontologies. Journal of Biomedical Informatics 39, 314–320 (2006). URL https://linkinghub.elsevier.com/retrieve/pii/S1532046406000190.

[27] Chen, Z. et al. A knowledge graph of clinical trials ($$\ mathop {\mathtt {CTKG }}\limits$$). Scientific Reports 12, 4724 (2022). URL https://doi.org/10.1038/s41598-022-08454-z https://www.nature.com/articles/s41598-022-08454-z.

[28] Reese, J. T. et al. KG-COVID-19: A Framework to Produce Customized Knowledge Graphs for COVID-19 Response. Patterns 2, 100155 (2021). URL http://dx.doi.org/10.1016/j.patter.2020.100155.

[29] Sanou, G. et al. IMGT-KG: A Knowledge Graph for Immunogenetics. Lecture Notes in Computer Science (including subseries Lecture Notes in Artificial Intelligence and Lecture Notes in Bioinformatics) 13489 LNCS, 628–642 (2022).

[30] Alshahrani, M., Thafar, M. A. & Essack, M. Application and evaluation of knowledge graph embeddings in biomedical data. PeerJ Computer Science 7, 1–28 (2021).

[31] Nicholson, D. N. & Greene, C. S. Constructing knowledge graphs and their biomedical applications. Computational and Structural Biotechnology Journal 18, 1414–1428 (2020). URL https://doi.org/10.1016/j.csbj.2020.05.017.

[32] Nguyen, D. Q. A survey of embedding models of entities and relationships for knowledge graph completion. arXiv (2017). URL http://arxiv.org/abs/1703.08098.

[33] Wang, Q., Mao, Z., Wang, B. & Guo, L. Knowledge Graph Embedding: A Survey of Approaches and Applications. IEEE Transactions on Knowledge and Data Engineering 29, 2724–2743 (2017). URL http://www.ieee.org/publications_standards/publications/rights/index.html http://ieeexplore.ieee.org/document/8047276/.

[34] Mohamed, S. K., Muñoz, E. & Novacek, V. On Training Knowledge Graph Embedding Models. Information 12, 147 (2021). URL https://www.mdpi.com/2078-2489/12/4/147/htm https://www.mdpi.com/2078-2489/12/4/147.

[35] Cambon M. and Cherouali K. and Kushwaha A. and Giudicelli V. and Duroux P. and Kossida S. and Lefranc M.-P. IMGT/mAb-DB and IMGT/2Dstructure-DB for IMGT standard definition of an antibody: from receptor to amino acid changes. Journées Ouvertes de Biologie Informatique et de Mathématiques (JOBIM) A597, 614–617 (2018). URL https://imgt.org/IMGTposters/522_Cambon_JOBIM2018_MPL_270718.pdf.

[36] Ferrara, N. & Adamis, A. P. Ten years of anti-vascular endothelial growth factor therapy. Nature reviews. Drug discovery 15, 385–403 (2016). URL https://pubmed.ncbi.nlm.nih.gov/26775688/.

[37] Ali, M. et al. PyKEEN 1.0: A Python Library for Training and Evaluating Knowledge Graph Embeddings (2020). URL http://arxiv.org/abs/2007.14175.

[38] Ali, M. et al. Bringing Light Into the Dark: A Large-scale Evaluation of Knowledge Graph Embedding Models Under a Unified Framework (2020). URL http://arxiv.org/abs/2006.13365.

[39] Abboud, R., Ceylan, I. I., Lukasiewicz, T. & Salvatori, T. BoxE: A box embedding model for knowledge base completion. Advances in Neural Information Processing Systems 2020-Decem, 1–13 (2020).

[40] Van Der Maaten, L. & Hinton, G. Visualizing Data using t-SNE. Journal of Machine Learning Research 9, 2579–2605 (2008).

[41] Rajabi, E. & Kafaie, S. Knowledge Graphs and Explainable AI in Healthcare (2022).

[42] Maćkiewicz, A. & Ratajczak, W. Principal components analysis (PCA). Computers & Geosciences 19, 303–342 (1993).

[43] Amid, E. & Warmuth, M. K. TriMap: Large-scale Dimensionality Reduction Using Triplets URL https://scikit-learn.org/stable/modules/generated/sklearn.datasets.make_s_curve.html.

[44] Reid, B. M., Permuth, J. B. & Sellers, T. A. Epidemiology of ovarian cancer: a review. Cancer Biology & Medicine 14, 9–32 (2017). URL https://www.cancerbiomed.org/content/14/1/9https://www.cancerbiomed.org/content/14/1/9.abstract.

[45] Matulonis, U. A. et al. Ovarian cancer. Nature Reviews Disease Primers 2, 1–22 (2016).

[46] Mabuchi, S., Morishige, K. & Kimura, T. Use of monoclonal antibodies in the treatment of ovarian cancer. Current opinion in obstetrics & gynecology 22, 3–8 (2010).

[47] Marmé, F. et al. Atezolizumab versus placebo in combination with bevacizumab and non-platinum-based chemotherapy in recurrent ovarian cancer: Final overall and progression-free survival results from the AGO-OVAR 2.29/ENGOT-ov34 study. Journal of Clinical Oncology 42, LBA5501–LBA5501 (2024). URL https://ascopubs.org/doi/10.1200/JCO.2024.42.17_suppl.LBA5501.

[48] Gonzàlez-Martín, A. et al. Atezolizumab Combined with Platinum and Maintenance Niraparib for Recurrent Ovarian Cancer with a Platinum-Free Interval ¿6 Months: ENGOT-OV41/GEICO 69-O/ANITA Phase III Trial. Journal of Clinical Oncology (2024). URL https://ascopubs.org/doi/10.1200/JCO.24.00668.

[49] Moore, K. N. et al. Atezolizumab, Bevacizumab, and Chemotherapy for Newly Diagnosed Stage III or IV Ovarian Cancer: Placebo-Controlled Randomized Phase III Trial (IMagyn050/GOG 3015/ENGOT-OV39). Journal of clinical oncology: official journal of the American Society of Clinical Oncology 39, 1842–1855 (2021). URL https://pubmed.ncbi.nlm.nih.gov/33891472/.

[50] Iril, C., Ozman, R., Milio, E. & Ontserrat, M. Chronic Lymphocytic Leukemia. New England Journal of Medicine 333, 1052–1057 (1995). URL https://www.nejm.org/doi/full/10.1056/NEJM199510193331606.

[51] Jaglowski, S. M., Alinari, L., Lapalombella, R., Muthusamy, N. & Byrd, J. C. The clinical application of monoclonal antibodies in chronic lymphocytic leukemia. Blood 116, 3705–3714 (2010). URL https://dx.doi.org/10.1182/blood-2010-04-001230.

[52] Shirley, M. Glofitamab: First Approval. Drugs 83, 1 (2023). URL https://pmc.ncbi.nlm.nih.gov/articles/PMC10245362/.

[53] Calabretta, E., Hamadani, M., Zinzani, P. L., Caimi, P. & Carlo-Stella, C. The antibody-drug conjugate loncastuximab tesirine for the treatment of diffuse large B-cell lymphoma. Blood 140, 303 (2022). URL https://pmc.ncbi.nlm.nih.gov/articles/PMC9335500/.

[54] Jin, S. et al. Emerging new therapeutic antibody derivatives for cancer treatment. Signal Transduction and Targeted Therapy 7, 1–10 (2022).

[55] Galkin, M., Yuan, X., Mostafa, H., Tang, J. & Zhu, Z. TOWARDS FOUNDATION MODELS FOR KNOWLEDGE GRAPH REASONING URL https://github.com/DeepGraphLearning/ULTRA.

[56] Cui, Y., Sun, Z. & Hu, W. A Prompt-Based Knowledge Graph Foundation Model for Universal In-Context Reasoning (2024). URL https://arxiv.org/abs/2410.12288v1.

